# Segmentation of Myelin-like Signals on Clinical MR Images for Age Estimation in Preterm Infants

**DOI:** 10.1101/357749

**Authors:** Maria Deprez, Siying Wang, Christian Ledig, Joseph V. Hajnal, Serena J. Counsell, Julia A. Schnabel

## Abstract

Myelination is considered to be an important developmental process during human brain maturation and closely correlated with gestational age. Assessment of the myelination status requires dedicated imaging, but the conventional T_2_-weighted scans routinely acquired during clinical imaging of neonates carry signatures that are thought to be associated with myelination. In this work, we propose a new segmentation method for myelin-like signals on T_2_-weighted magnetic resonance images that could be used to assess neonatal brain maturation in clinical practice. Firstly we define a segmentation protocol for myelin-like signals, and delineate manual annotations according to this protocol. We then develop an expectation-maximization framework through which we obtain the automatic segmentations of myelin-like signals. We incorporate an explicit class for partial volume voxels whose locations are configured in relation to the composing pure tissues via second-order Markov random fields. We conduct experiments in the thalami and brainstem where the majority of myelination occurs during the perinatal period for 16 test subjects aged between 29 and 44 gestational weeks. The proposed method performs accurately and robustly in both regions with respect to the manual annotations over a range of intensity percentile thresholds that are used to generate the initial segmentation estimates. Finally, we construct spatio-temporal growth models for myelin-like signals in the thalami and brain-stem to demonstrate the applicability of the proposed method for age estimation in preterm infants.

## 1. Introduction

Human brain maturation involves a complex series of morphological, structural and functional changes. Among these changes is the process of myelin growth and axon ensheathment known as myelination, which facilitates electrical conduction in the neural system [19]. In clinical practice, birth weights are commonly used for measuring gestational maturity [26]. However, Rorke *et al.* [26] reported that some well-developed infants based on the measurements of body weight show retarded brain maturation, whereas the neurodevelopmental patterns in preterm infants with extensive myelination are similar to those found in fullterm babies, regardless of the birth weights. Considering the critical role of myelination in neurodevelopment [12], it is of great interest to utilize myelin as an early marker of abnormal brain development in newborns.

Non-invasive non-ionizing magnetic resonance imaging (MRI) can be used to track myelination in the developing brain, for example T_1_-weighted (T_1_w) and T_2_w imaging, myelin water imaging (MWI) [1], magnetization transfer imaging (MTI) [16], and diffusion tensor imaging (DTI) [20]. In this work, we use T_2_w images, where myelin appears as low intensities, due to the close correlation between T_2_ relaxation time and the final stage of myelination [11]. Moreover, T_1_w and T_2_w acquisitions remain the most routine MRI techniques compared to the advanced MWI, MTI and DTI which require elaborate scans that are constrained in clinical practice when imaging fragile preterm neonates [29, 32]. Qualitative descriptions of myelination on T_2_w images [4, 8] are in great consistency with histological observations [42]. Thus the ability to numerically assess myelination using T_2_w scans would facilitate neurodevelopmental evaluation in preterm infants directly from large-scale clinical MRI data.

In this study, we refer to the tissue that is likely to contain myelin in T_2_w neonatal brain MRI as “myelin-like signals” (MLS). Automatic segmentation of MLS is challenging. Firstly, myelin is not included in any of the existing neonatal brain atlases [18, 30] or manual annotation database [13]. Most neonatal brain segmentation methods adapt the approaches developed for adults [3, 15, 34, 39, 43], and use a probabilistic atlas or manual annotations to obtain prior information on the expected tissue locations. Prastawa *et al.* [25] simulated a newborn brain atlas by averaging the semi-automatic segmentations from three subjects, and separated the myelinated and unmyelinated white matter (WM) using minimum spanning tree. Xue *et al.* [41] performed intensity-based *k*-means clustering that divided the developing brain into gray matter (GM), WM and cerebrospinal fluid (CSF) without requiring any probabilistic or manual atlases. Anbeek *et al.* [2] trained a *k*NN classifier using samples selected from the manual annotations of 12 patients based on the features of voxel intensities and spatial locations. Recently developed methods in the NeoBrainS12 challenge [14, 22, 24, 38, 40] all showed great promise in general neonatal brain segmentation. However, none of them performed well in segmenting myelination due to insufficient spatial prior information [17]. Secondly, partial volume (PV) voxels containing both MLS and the background (BKG) tissue in a region of interest (ROI) need to be modeled explicitly when segmenting MLS. Fig. 1 illustrates that a simple two-class Gaussian mixture model (GMM) without PV modeling fails to separate the intensity distributions of MLS and BKG. The current PV estimation methods [6, 21, 23, 31, 33, 36] depend on prior knowledge of the spatial locations of the composing pure tissues to guide the search for PV voxels, which is, however, very limited in MLS segmentation.

**Figure 1:**
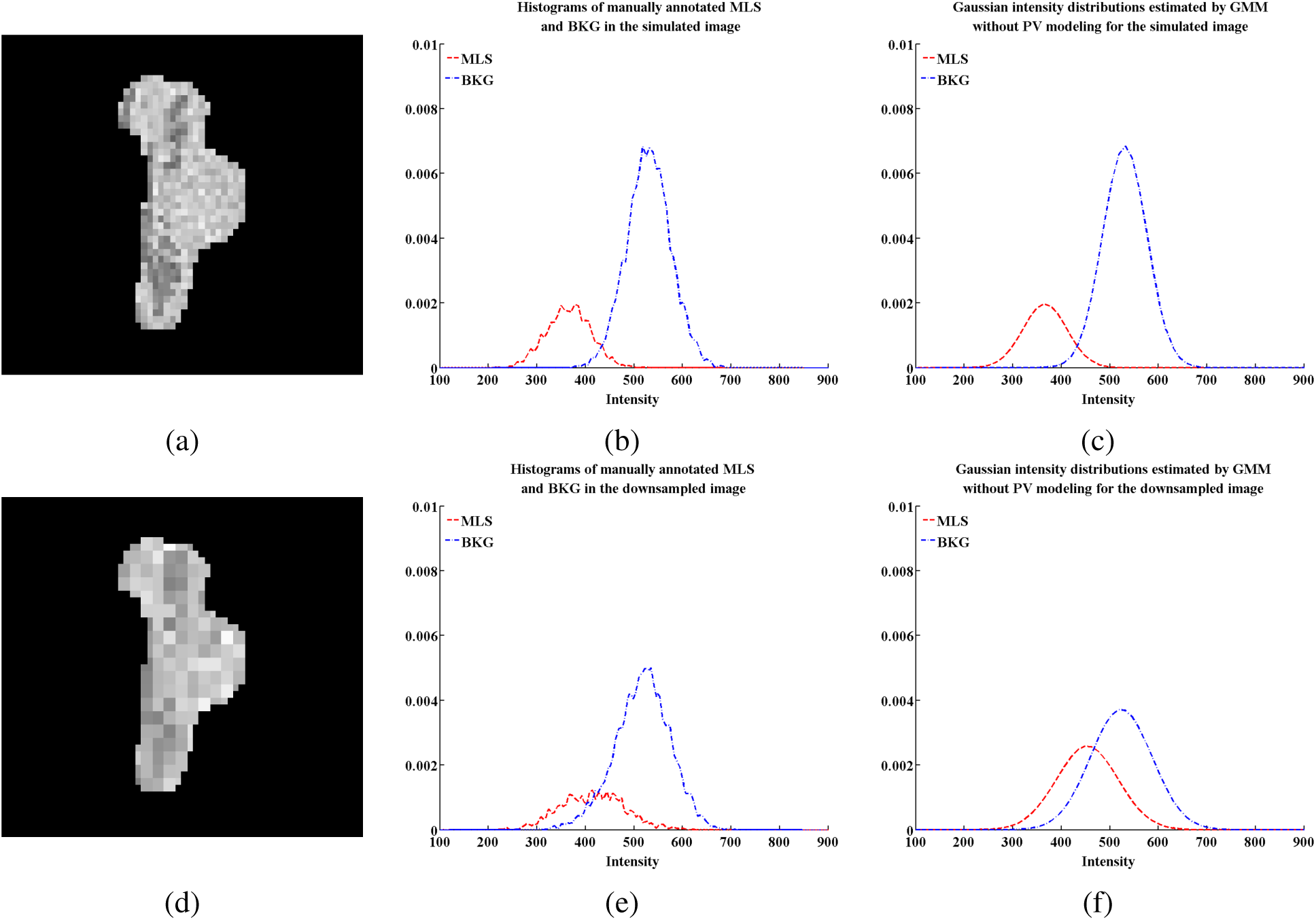
Simulated T_2_-weighted images in the brainstem (sagittal view) for a preterm infant at 38 gestational weeks (a) before and (d) after downsampling. The intensities of myelin-like signals (MLS) and the remaining background (BKG) are generated from two Gaussians, and the MLS voxels are labeled based on the manual annotation. A two-class Gaussian mixture model (GMM) recovers the distributions correctly before downsampling. Partial volume (PV) voxels created after downsampling shift the histogram of manually annotated MLS towards BKG, and the simple GMM without PV estimation can no longer approximate the histograms accurately.

We therefore develop a new expectation-maximization (EM) framework for MLS segmentation on T_2_w neonatal brain images that does not require any probabilistic atlas or manual annotation of myelin. We introduce an explicit PV class whose locations are configured in relation to MLS and BKG in a predefined ROI using a 3D connectivity tensor via second-order Markov random fields (MRFs) [21]. This approach allows us to distinguish the small proportion of MLS in the presence of substantial PV voxels. Our method achieves automatic MLS segmentations of high Dice coefficients (DCs) [10] with respect to the manual annotations for 16 preterm infants at one-week intervals between 29 and 44 weeks gestational age (GA). We further use the segmentations obtained from 114 preterm infants to build a spatio-temporal model of progressing myelination.

This paper is structured as follows. In Section 2, we define an MLS segmentation protocol in two ROIs, namely the thalami and brainstem. Manual annotations are delineated according to this protocol. Section 3 presents the proposed MLS segmentation method, followed by the experiments and results in Section 4. Finally, we demonstrate the applicability of this segmentation approach for assessing neonatal brain development in Section 5.

## 2. Materials

### 2.1. Subjects and MR acquisition

114 preterm infants without brain injuries were scanned between 29 and 44 weeks GA at Hammersmith Hospital, London, UK. Ethical permission for this study was granted by the Hammersmith and Queen Charlotte’s and Chelsea Research Ethics Committee (07/H0707/101). Written parental consent was obtained prior to scanning. T_2_w fast spin-echo brain images were acquired on a 3T Philips Intera system with repetition time = 8700 ms, echo time = 160 ms and voxel sizes = 0.86 mm*×*0.86 mm*×*1 mm.

### 2.2. Segmentation protocol and manual annotations

We choose the thalami and brainstem as the ROIs for assessing MLS because these are the regions where the majority of myelination occurs between 29 and 44 weeks GA [8, 28]. The automatic procedure through which we obtain the binary ROI masks of the thalami and brainstem for individual subjects is explained in Section 3.1.

The MLS segmentation protocol is defined within the ROIs based on both image intensities and anatomical knowledge [8]. We delineate three central brain structures in the thalami, including the posterior limbs of the internal capsule (PLIC), ventrolateral nuclei (VLN) and subthalamic nuclei (STN). Particularly the PLIC tracts in the preterm brain, which show MLS at approximately 40 weeks GA on T_2_w images [8], is an important signature that can be used to assess neonatal brain maturation [5]. We define the segmentation protocol in the brainstem mainly based on intensities because the image resolution is insufficient to identify the detailed anatomical structures.

Manual annotations of MLS, delineated in the ROIs according to the defined segmentation protocol, are available for 16 preterm infants, each at a different time point, between 29 and 44 weeks GA at one-week intervals. An example is shown in Fig. 2. Eight of the 16 subjects, from 30 weeks GA onward at two-week intervals, have a repeated manual segmentation delineated by the same rater in order to assess the intra-rater reliability. The quality of these manual annotations was confirmed by a clinical expert.

**Figure 2:**
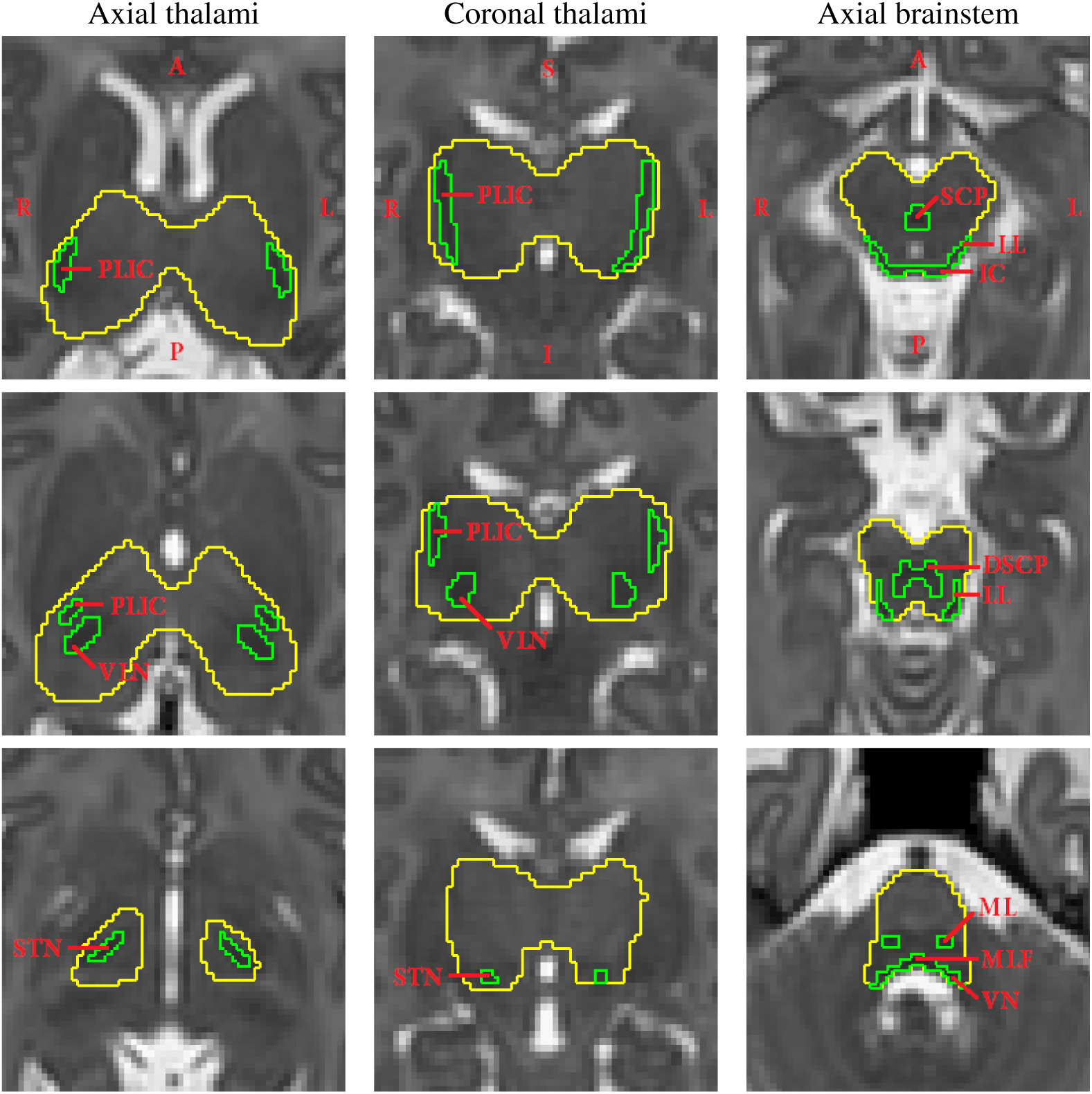
Manual annotations of myelin-like signals (green label) in the thalami and brainstem of a subject at 42 gestational weeks, delineated according to the defined segmentation protocol. The regions of interest are labeled in yellow. The columns from left to right show the thalami in the axial and coronal views, and the brainstem in the axial view. Abbreviations: PLIC–posterior limb of the internal capsule, VLN–ventrolateral nucleus, STN–subthalamic nucleus, SCP–superior cerebellar peduncle, DSCP–decussation of the superior cerebellar peduncle, IC–inferior colliculus, LL– lateral lemniscus, ML–medial lemniscus, MLF–medial longitudinal fasciculus, VN–vestibular nucleus.

## 3. Method

The overall pipeline proposed for MLS segmentation on T_2_w neonatal brain MR images is shown in Fig. 3. In this section, we first explain the procedure of image preprocessing. We then describe a purely intensity-based GMM without PV modeling, followed by the incorporation of an explicit PV class and spatial regularization in the EM framework. Lastly, we explain the implementation details of the proposed method for obtaining the final binary MLS segmentations.

**Figure 3:**
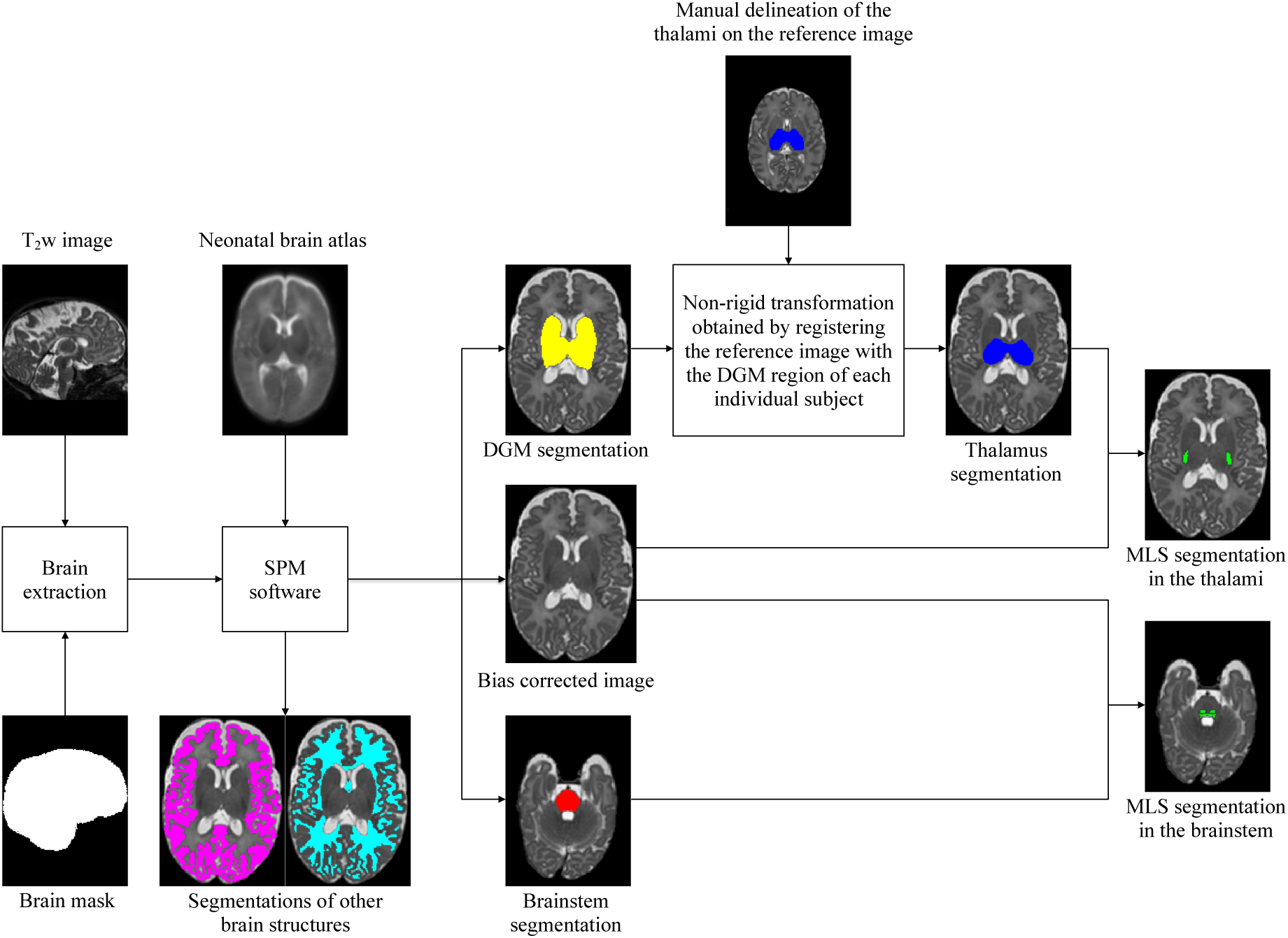
Schematic flow of the proposed segmentation approach for myelin-like signals (MLS) on T_2_w neonatal brain images. Through image preprocessing, we achieve brain extraction and bias field correction, and obtain the binary masks of the thalami and brainstem for individual subjects. The automatic segmentations of the deep gray matter (DGM), obtained using the Statistical Parametric Mapping (SPM) software, contain the basal ganglia, thalami, hippocampi and amygdalae. We extract the masks for the thalami from the DGM segmentations via label propagation.

### 3.1. Image preprocessing

The aim of image preprocessing is to achieve brain extraction and bias field correction, and to obtain the binary ROI masks of the thalami and brainstem for individual subjects.

We remove the non-brain tissues using label propagation [15] of manually annotated brain masks, and segment the skull-stripped images using the Statistical Parametric Mapping (SPM) software^2^ (version SPM8) [3]. The spatial priors are provided by a 4D probabilistic neonatal brain atlas^3^ [18]. Subsequently, we obtain the bias-corrected images as well as the segmentations of the cortical GM, WM, deep gray matter (DGM), brainstem, cerebellum and CSF.

We use the automatic SPM segmentations of the brainstem directly as the ROI masks. The DGM segmentations include the basal ganglia, thalami, hippocampi and amygdalae. Since the myelinated thalamic nuclei can have similar intensities to the densely organized GM in the basal ganglia which does not contain myelin between 29 and 44 weeks GA [8], we extract the ROI masks for the thalami from the DGM segmentations in order to prevent misclassifications by the intensity-based segmentation model. We first manually delineate the thalami on the T_2_w image of a single reference subject at 36 weeks GA. We register the reference image with the dilated DGM region of each subject using free-form deformation (FFD) non-rigid registration [27], and then transform the manual delineation of the thalami from the reference space to each individual subject’s space. The ROI masks of the thalami and brainstem are illustrated in Fig. 4.

**Figure 4:**
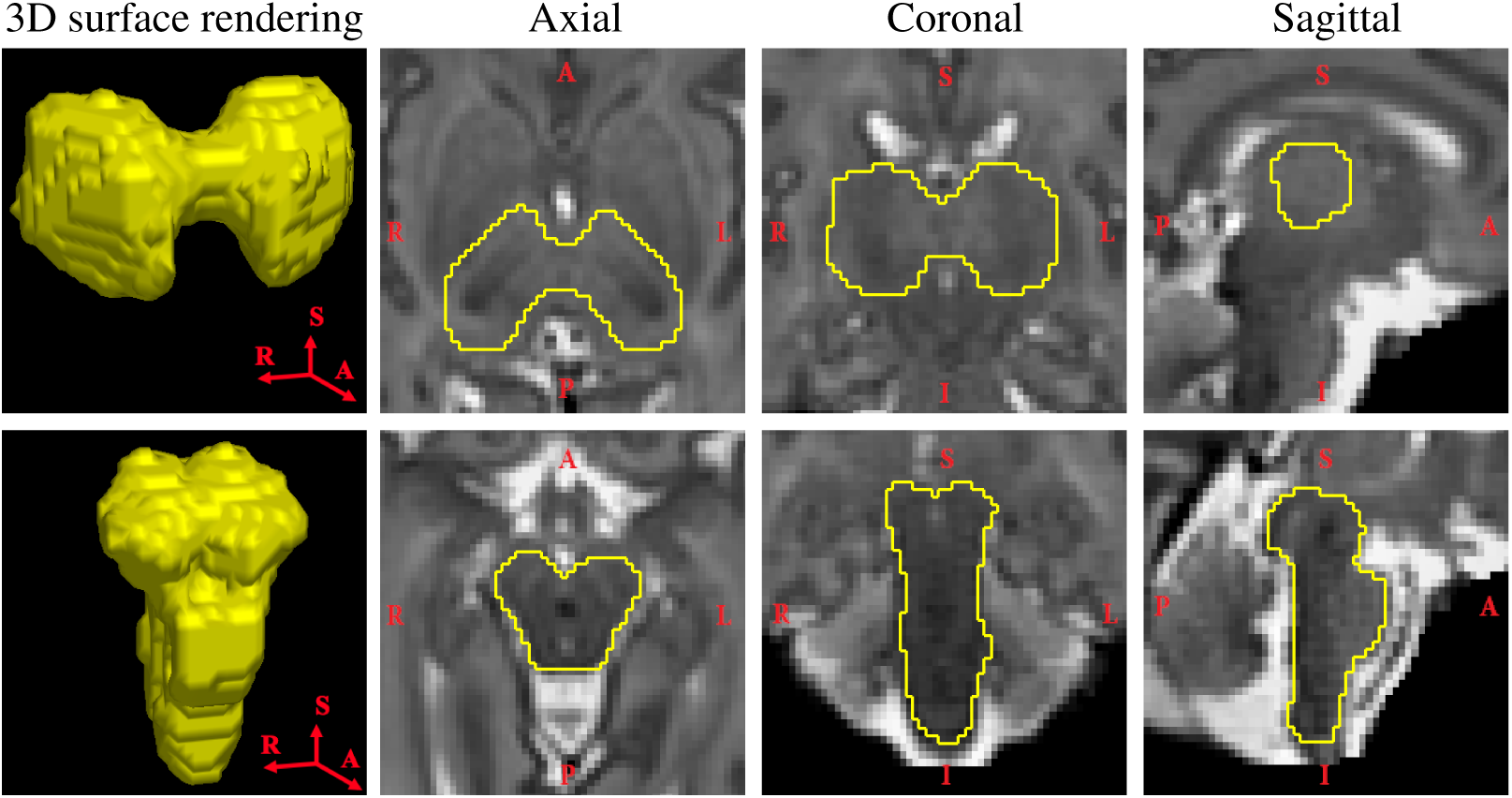
Two separate regions of interest (ROIs) labeled in yellow. The first and second rows show the thalami and brainstem, respectively, for a subject at 34 gestational weeks. The columns from left to right show the ROIs in 3D surface rendering, axial, coronal and sagittal views.

### 3.2. Purely intensity-based GMM without PV modeling

First we segment MLS using a standard GMM without PV modeling. We define two classes in the ROI: myelin-like signals (MLS) and background (BKG). The conditional probability density function (PDF) of class *k* is approximated as a Gaussian with mean *μ_k_* and standard deviation (SD) *σ_k_*. The prior probability that voxel *i* belongs to class *k* is modeled by the spatially constant mixing proportion *c_k_*. We apply the EM algorithm [9] as follows:

- E-step: Calculate the probability *p_ik_* that voxel *i* belongs to class *k*:

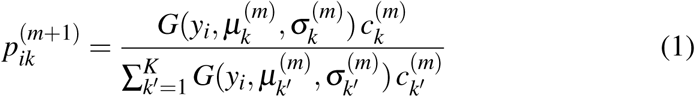

where *y_i_* is the observed intensity, *K* the number of classes, and *m* the current EM iteration number.
- M-step: We assume that the SDs of the MLS and BKG classes are equal. Ideally the MLS and BKG tissues in the ROI have uniform intensities. Variations arise mainly due to noise and can be considered constant across the MR image. Therefore, the two classes may be regarded as Gaussians with equal SDs provided that their mean intensities are distant from zero. Based on this assumption, the maximum likelihood (ML) estimations for the GMM parameters of class *k* yield the following equations:

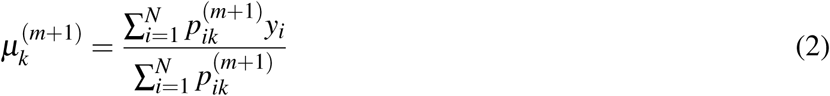

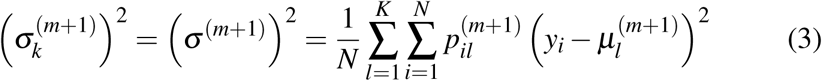

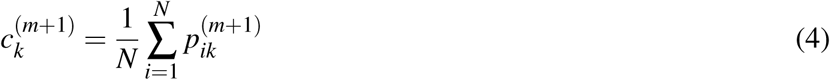

where *N* is the number of voxels in the ROI.

### 3.3. Explicit PV modeling

As explained in Fig. 1, MLS becomes much less distinguishable in the histogram of the downsampled image due to the intermediate PV intensities. The simple two-class GMM described in Section 3.2 over-estimates the MLS class and cannot recover the Gaussians before downsampling. Therefore, we introduce an explicit PV class in addition to the previous MLS and BKG classes to model the intensities of mixed tissues.

The E-step (Eq. 1) remains unchanged at this stage. In the M-step, we approximate the PV class mean as the arithmetic mean of the composing pure tissue means, and assume that the SDs of the MLS, PV and BKG classes are identical for the reason explained in Section 3.2:

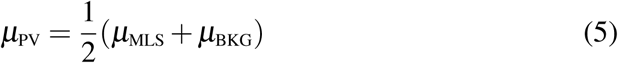

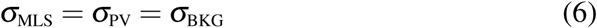

By substituting Eqs. 5 and 6 into the objective function of log likelihood [35], the SD *σ_k_* of class *k* can be updated using Eq. 3 and the ML estimations for the means of the MLS and BKG classes result in the following system of linear equations (see more details in the appendix):

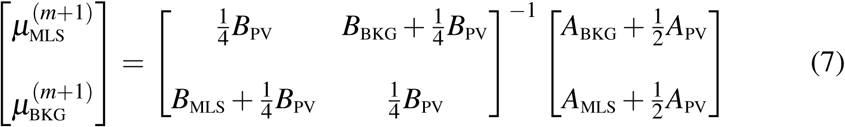

where

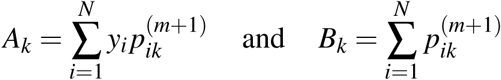

### 3.4. Spatial regularization via second-order MRFs

Van Leemput *et al.* [36] pointed out that introducing an additional PV class without any spatial prior can disrupt the process of EM convergence. Therefore, robust PV estimation methods generally require prior knowledge of the spatial locations of the composing pure tissues to guide the search for PV voxels [6, 21, 23, 31, 33, 36]. However, MLS is not included in any of the existing neonatal brain atlases [18, 30] or manual annotation database [13]. To overcome this difficulty, we incorporate spatial regularization in the EM framework via secondorder MRFs [21] where we configure the dependencies among a triplet of classes in the neighborhood using a 3D connectivity tensor (Fig. 5).

**Figure 5:**
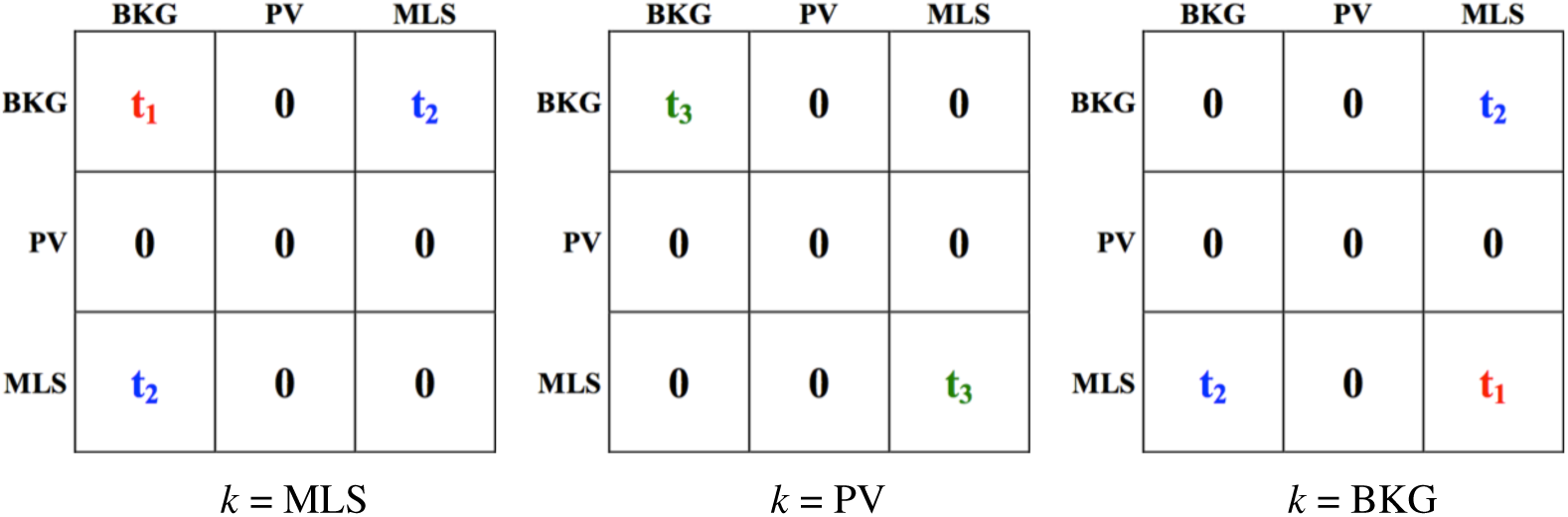
Matrix **T***_k_* in the 3D connectivity tensor for *k* = MLS, PV and BKG. Each ele ment **T***_k_*(*k*_1_*, k*_2_) indicates the penalty when classes *k*_1_ and *k*_2_ are both present in the neighborhood of class *k*. We will determine the values of penalties *t*_1_, *t*_2_ and *t*_3_ empirically in Section 4.1. Abbreviations: MLS–myelin-like signals, PV–partial volume, BKG–background.

First we modify the E-step (Eq. 1) by replacing the spatially constant mixing proportion *c_k_* with the spatial prior 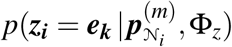 acquired from the neighborhood as described in [35]:

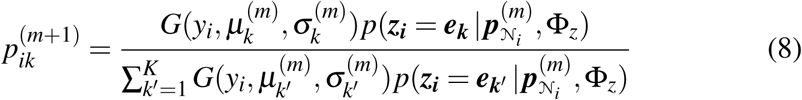

where

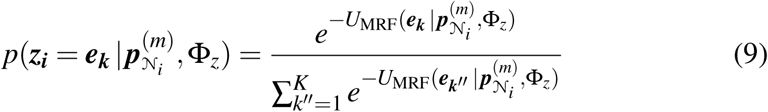

The symbol ***z_i_*** indicates to which class voxel *i* belongs, and ***e_k_*** is a unit vector of length *K* whose components are equal to zero except the *k*th component. The probabilities in the 26-neighborhood 𝒩*_i_* of voxel *i* are denoted as ***p***_𝒩_*i*__, and the MRF parameters as Φ*_z_*. We compute the energy function *U*_MRF_(***e_k_*** _|_ ***p***_𝒩_*i*__, Φ*_z_*) for second-order MRFs as follows:

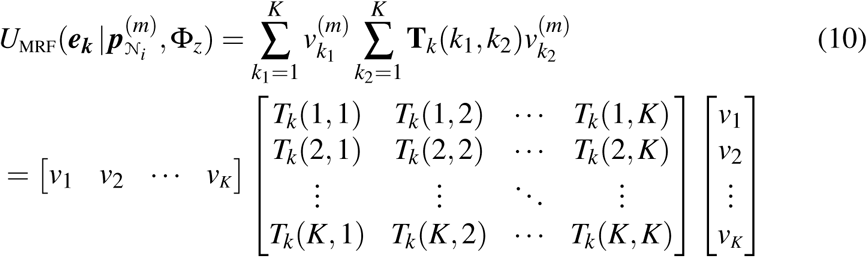

where

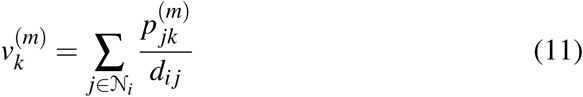

Here *p_jk_* is the probability that neighbor *j* belongs to class *k*, and *d_i j_* the Euclidean distance between voxel *i* and neighbor *j*. The element of the connectivity tensor, denoted as **T***_k_*(*k*_1_*, k*_2_), indicates the penalty when classes *k*_1_ and *k*_2_ are both present in the neighborhood of class *k*. By assigning a large value to **T***_k_*(*k*_1_*, k*_2_) in Eq. 10, we are able to penalize a particular configuration when the probabilities of classes *k*_1_ and *k*_2_ are both high in the neighborhood. This would result in a large value of the energy function *U*_MRF_(***e_k_*** | ***p***_𝒩_*i*__, Φ*_z_*), making voxel *i* less probable to belong to class *k*.

We assign the penalties **T***_k_*(*k*_1_*, k*_2_) according to the following rules based on the triplet that contains class *k* of the current voxel and classes *k*_1_ and *k*_2_ of a pair of neighboring voxels:

- **T***_k_*(MLS, MLS) and **T***_k_*(BKG, BKG): We set the penalty to 0 if class *k* is the same as the pair. We assign a penalty *t*_1_ if *k* is a pure tissue class different from the pair to encourage the PV class as MLS and BKG are both in the triplet. We assign a penalty *t*_3_ if *k* is PV as there is no evidence for PV in the triplet. This effectively prevents over-estimation of the PV class.
- **T***_k_*(MLS, BKG): We assign a penalty *t*_2_ if *k* is MLS or BKG, and set the penalty to 0 if *k* is PV. We aim to encourage the PV class by deliberately penalizing the MLS and BKG classes if they are both present in the neighborhood.
- **T***_k_*(MLS, PV), **T***_k_*(PV, PV) and **T***_k_*(BKG, PV): We set the penalty to 0 regardless of the class of *k*.

The values of penalties *t*_1_, *t*_2_ and *t*_3_ will be determined empirically in Section 4.1. We again assume that the PV class mean equals the arithmetic mean of the composing pure tissue means and that the SDs of the MLS, PV and BKG classes are identical. Provided that the penalties remain constant, the SD *σ_k_* and mean *μ_k_* of class *k* in the EM framework with MRFs can be estimated using Eqs. 3 and 7, respectively, in the same way as the purely intensity-based GMM with an explicit PV class (see the appendix for details).

### 3.5. Implementation

The EM algorithm is initialized as follows. We first sort the T_2_w intensities in the thalami of individual subjects in ascending order and choose the 6th percentile as the threshold. This is because the average volume fraction of manually annotated MLS in the thalami is 6% over the 16 test subjects at one-week intervals between 29 and 44 weeks GA. Voxels with intensities below the threshold constitute the initial MLS segmentation, and the remaining voxels in the ROI form the initial BKG segmentation. We will verify in Section 4.3 that the percentile estimated this way optimizes the MLS segmentations that could possibly be achieved for all the 16 subjects using a single threshold. We assign the initial MLS and BKG segmentations to the corresponding posterior probability maps (PPMs), and set the PPM of the PV class equal to zero. Similarly, we threshold the T_2_w intensities in the brainstem of individual subjects at the 25th percentile to obtain the initial segmentation estimates in this region.

Tissue classification is achieved by interleaving the E-step and M-step as described in Section 3.4 until the relative change in the objective function of log likelihood [35] is less than 0.01%. The resultant PPMs are converted into hard segmentations using the maximum-vote rule. In addition, we reclassify the PV voxels as one of the composing pure tissues by calculating the fraction of the MLS class, denoted as *f_i_*, at PV voxel *i*:

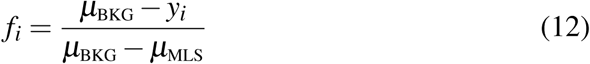

where *μ*_MLS_ and *μ*_BKG_ are the means of the MLS and BKG classes, and *y_i_* the observed intensity. PV voxels with fractions above 0.5 are reclassified as MLS and combined with the voted hard segmentation of the MLS class to form the final segmentation.

## 4. Experiments and results

In this section, we first determined the optimal values of the penalties in the connectivity tensor for the proposed method with explicit PV modeling via second-order MRFs (GMM-PV-MRF). We then evaluated GMM-PV-MRF in terms of Dice overlap with manually annotated MLS, as compared to two other segmentation models in Table 1. The initial MLS segmentations generated by optimal thresholding exploited manual annotations of MLS that are usually unavailable in practice. We thus further investigated the dependence of each method on the initial threshold. Lastly, we assessed the intra-rater reliability compared to the accuracies of the automated model-based methods.

**Table 1:** Segmentation models for myelin-like signals.

| Segmentation model | Description | Partial volume (PV) modeling | Spatial regularization |
| --- | --- | --- | --- |
| GMM | Purely intensity-based Gaussian mixture model without PV modeling | × | × |
| GMM-PV | GMM incorporating a spatially unconstrained PV class | ✓ | × |
| GMM-PV-MRF | GMM with explicit PV modeling via second-order MRFs | ✓ | ✓ |

### 4.1. Optimization of the penalties in the connectivity tensor

As explained in Section 3.4, we acquired the MLS, PV and BKG priors by configuring their spatial dependencies in the connectivity tensor (Fig. 5) via second-order MRFs. For each individual ROI, we optimized the penalties *t*_1_, *t*_2_ and *t*_3_ using a coarse-to-fine grid search method.

We first determined the coarse value of each penalty from the following set {0.003, 0.007, 0.03, 0.07, 0.3, 0.7}. Two values were included for the lower and upper halves of each tenfold scale. We computed MLS segmentations using all 216 combinations of the *t*_1_, *t*_2_ and *t*_3_ values. We chose the combination that yielded the highest average DC over the 16 test subjects with respect to the manual annotations. A DC is defined for two overlapping volumes as the ratio of the number of voxels in the intersection to the average number of voxels in the two volumes:

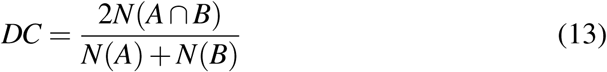

where *N*(*A*) denotes the number of voxels in volume *A*.

We then performed a fine grid search through ten different values for each penalty with the selected coarse value at the center. For instance, we examined the set {0.007, 0.008, 0.009, 0.01, 0.02, 0.03, 0.04, 0.05, 0.06, 0.07} for *t*_1_ in the thalami, whose best value at the coarse level was found to be 0.03. The test sets and optimal values of all three penalties at the coarse and fine levels are summarized in Table 2 for both ROIs. The final (*t*_1_*, t*_2_*, t*_3_) combination that yielded the highest average DC was (0.05, 0.03, 0.01) in the thalami, and (0.05, 0.03, 0.009) in the brainstem, which were highly consistent even though optimized separately.

**Table 2:**
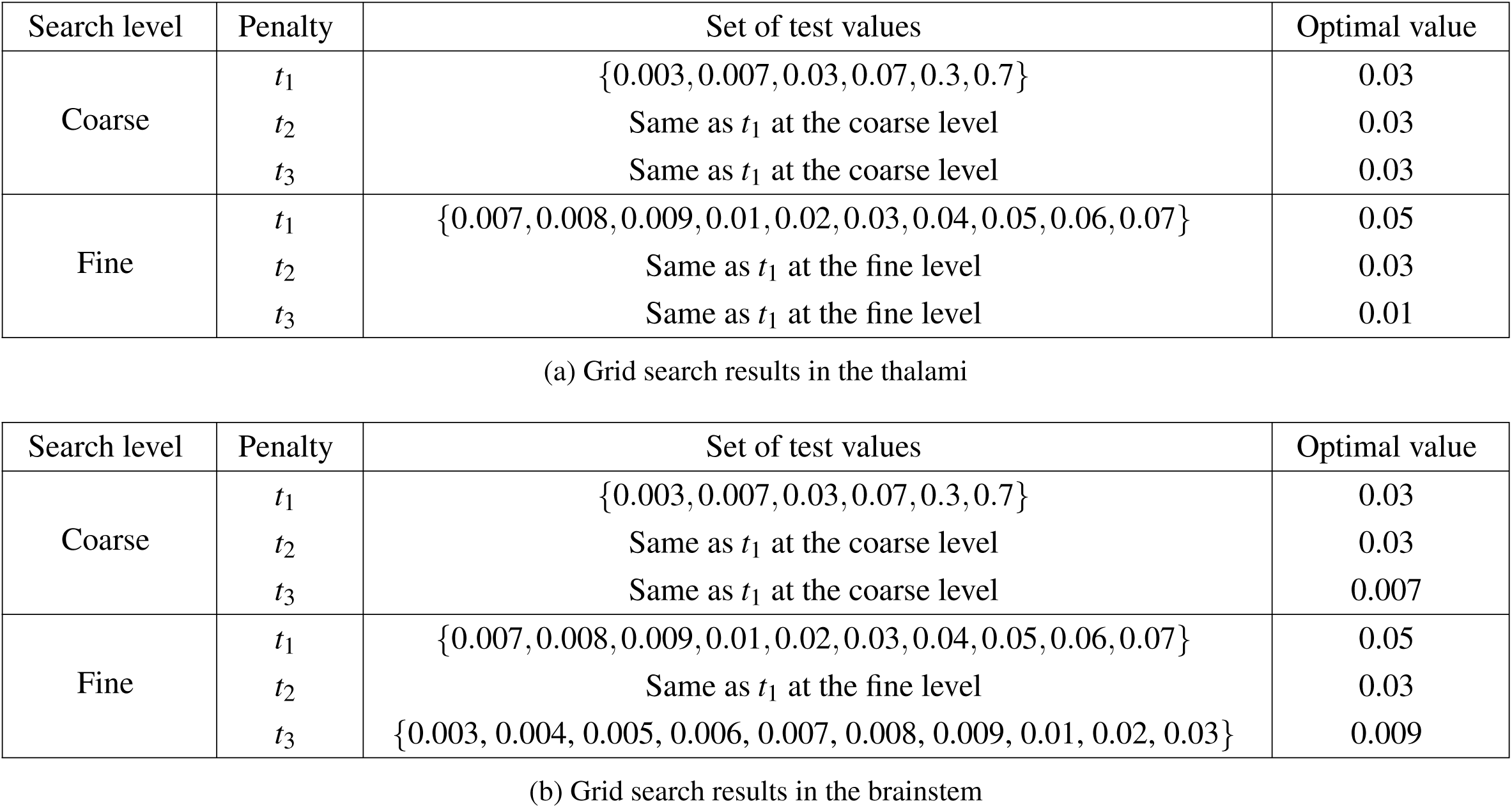
Optimal values of the penalties *t*_1_, *t*_2_ and *t*_3_ in different regions of interest (ROIs) determined using a coarse-to-fine grid search method. These penalties are assigned to the connectivity tensor (Fig. 5) for configuring spatial dependencies of myelin-like signals, partial volume voxels and the background in each ROI.

### 4.2. Evaluation of the MLS segmentation methods

We segmented MLS using GMM, GMM-PV and GMM-PV-MRF (see Table in the thalami and brainstem for the 16 test subjects at one-week intervals between 29 and 44 weeks GA. From here on, we used the optimal values of *t*_1_, *t*_2_ and *t*_3_ determined in Section 4.1 for GMM-PV-MRF. We implemented GMM and GMM-PV as described in Sections 3.2 and 3.3 respectively.

The DCs of the resultant segmentations with respect to the manual annotations are summarized for both ROIs in Table 3. We also evaluated the initial MLS segmentations obtained by optimal thresholding as a baseline method. We compared GMM-PV-MRF with all the other methods using two-tailed Student’s *t*-tests at the 5% significance level, and the *p*-values are shown in Table 3.

**Table 3:**
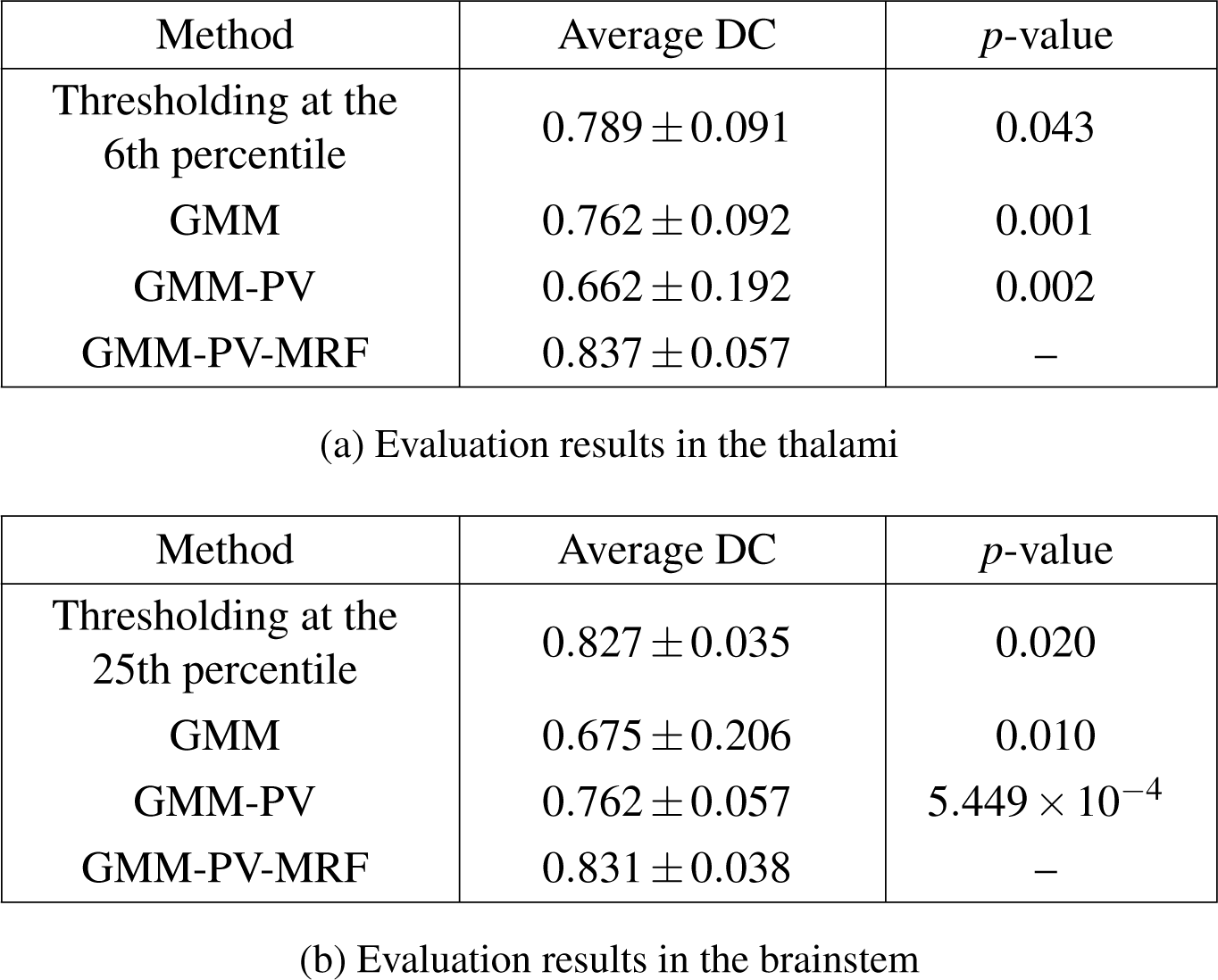
Average Dice coefficients (DCs) (*±* standard deviations) of myelin-like signals segmented by optimal thresholding and the model-based methods (see Table 1 for method descriptions) for 16 test subjects aged between 29 and 44 gestational weeks. GMM-PV-MRF is compared with all the other methods using two-tailed Student’s *t*-tests at the 5% significance level. GMM-PV-MRF achieves the highest average DC and significantly outperforms all the other methods in both regions of interest.

Thresholding at the optimal level resulted in a lower average DC and a larger SD in the thalami than in the brainstem. The 6th percentile threshold was derived from the average volume fraction of manually annotated MLS in the thalami over the 16 subjects, and hence decreased the DCs at the early and late GAs due to the appearance of new myelination in this region [8]. In contrast, the myelinated brainstem grew with the overall increase in the brain size with relatively little new myelination appearing between 29 and 44 weeks GA [7, 28], which implied an approximately constant MLS volume fraction.

The purely intensity-based GMM without PV modeling displayed large segmentation variations in the brainstem compared to the thalami. In Figs. 6 and 7, we studied the Gaussian distributions of the PPMs estimated using different segmentation models for a subject at 37 weeks GA. With respect to the manual annotation histograms, GMM over-estimated the MLS class in the thalami with a shift of the mean towards BKG, and failed to separate the two classes in the brainstem. This is caused by PV voxels represented as the overlapping area between the histograms of manually annotated MLS and BKG.

**Figure 6:**
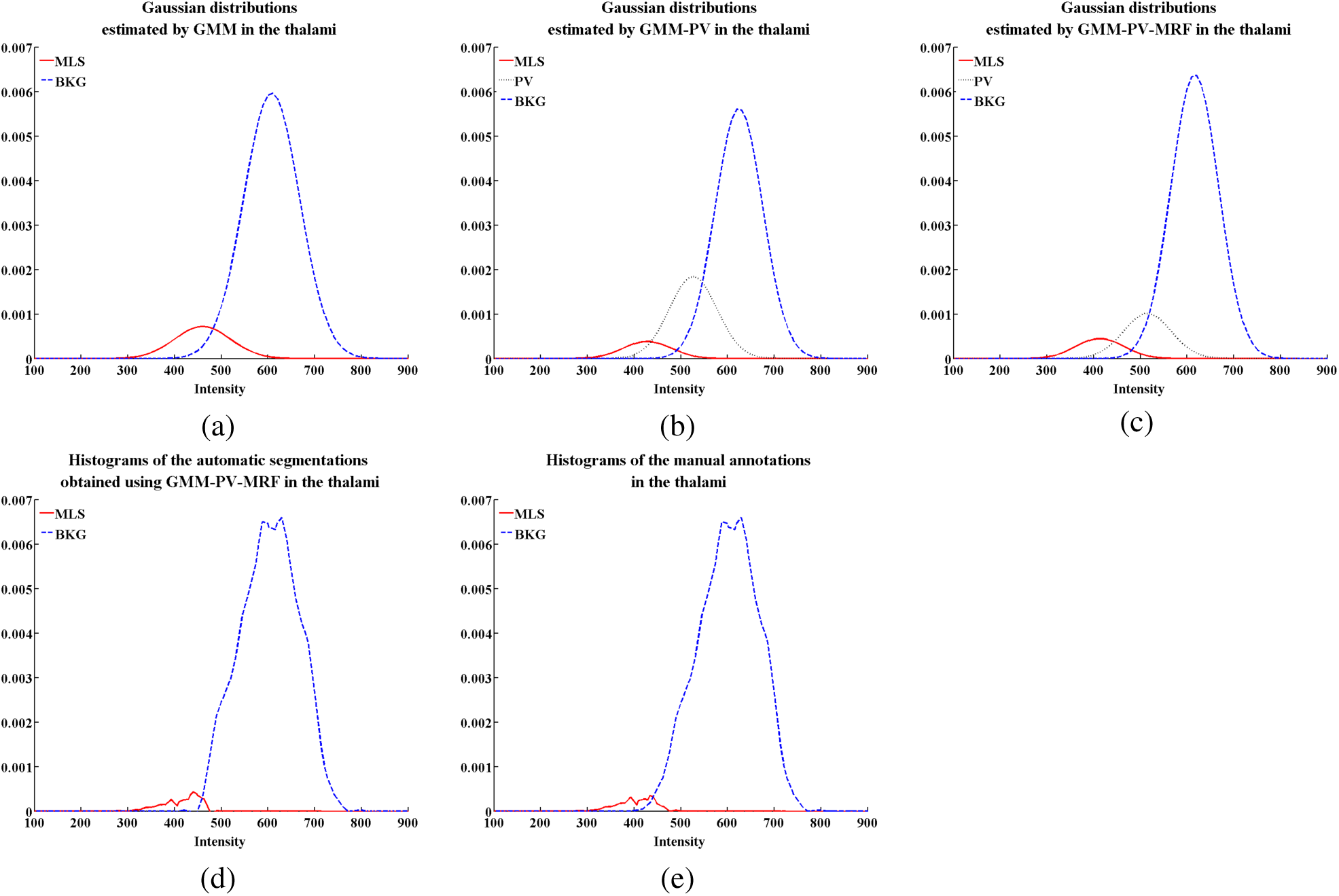
Gaussian distributions of the posterior probability maps estimated using the segmentation models in Table 1 for myelin-like signals (MLS), background (BKG) and partial volume voxels (PV) in the thalami of a subject at 37 gestational weeks. The sum of the graph areas for all the classes equals one. GMM over-estimates the MLS class with a shift of the mean towards BKG compared to the manual annotation histograms. GMM-PV over-estimates the PV class compared to GMM-PV-MRF due to the absence of spatial regularization, as confirmed in Fig. 8(c). The histograms of the binary segmentations obtained using GMM-PV-MRF are almost the same as those of the manual annotations.

**Figure 7:**
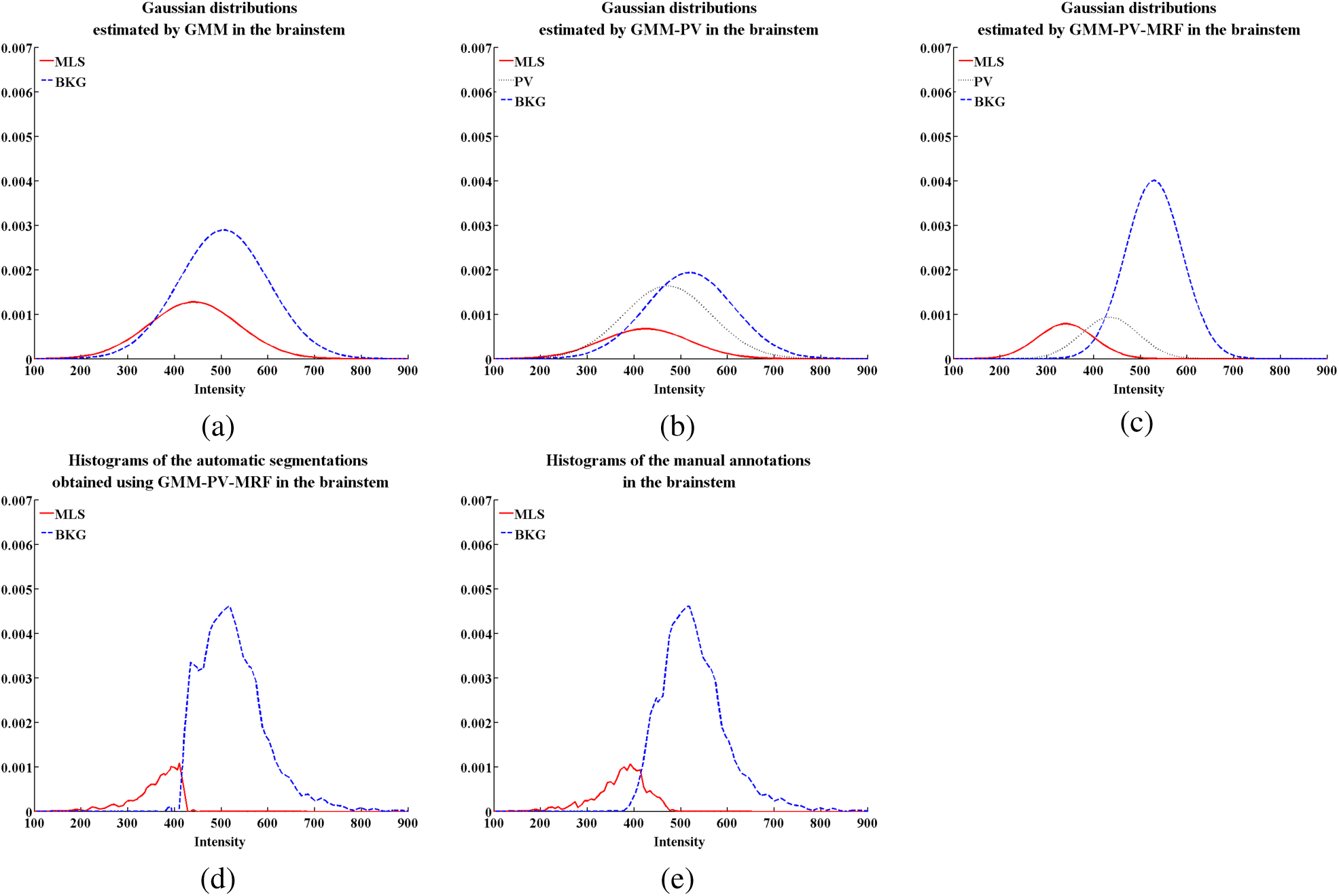
Gaussian distributions of the posterior probability maps estimated using the segmentation models in Table 1 for myelin-like signals (MLS), background (BKG) and partial volume voxels (PV) in the brainstem of a subject at 37 gestational weeks. The distributions and histograms are shown in the same scale as Fig. 6. The brainstem contains more PV voxels than the thalami, which can be seen from the larger overlapping area between the histograms of manually annotated MLS and BKG. EM classification only succeeds in GMM-PV-MRF, and the histograms of the resultant binary segmentations highly resemble those of the manual annotations.

The extra Gaussian in GMM-PV caused large segmentation variations in the thalami due to the absence of spatial regularization. In the brainstem, GMM-PV appeared to reduce the variations previously shown by GMM, but nonetheless yielded significantly lower DCs (*p <* 0.05) than the baseline method of optimal thresholding. Fig. 9(c) revealed that, without any spatial constraints, the PPM of the PV class in GMM-PV for the same subject at 37 weeks GA spread across the brainstem with little variation in space.

GMM-PV-MRF achieved high DCs consistently across all the GAs in both ROIs. The extra Gaussian modeled the intermediate PV intensities, and the spatial constraints imposed via second-order MRFs confined the PV locations at the boundary between MLS and BKG. This approach prevented the PV Gaussian distribution from shifting towards BKG and hence the spread of the PV class in space, as demonstrated in Figs. 6(c), 7(c), 8(f) and 9(f). The histograms of the binary segmentations obtained using GMM-PV-MRF in Figs. 6(d) and 7(d) highly resembled those of the manual annotations. Student’s *t*-tests in Table 3 confirmed that GMM-PV-MRF outperformed all the other methods in both ROIs.

**Figure 8:**
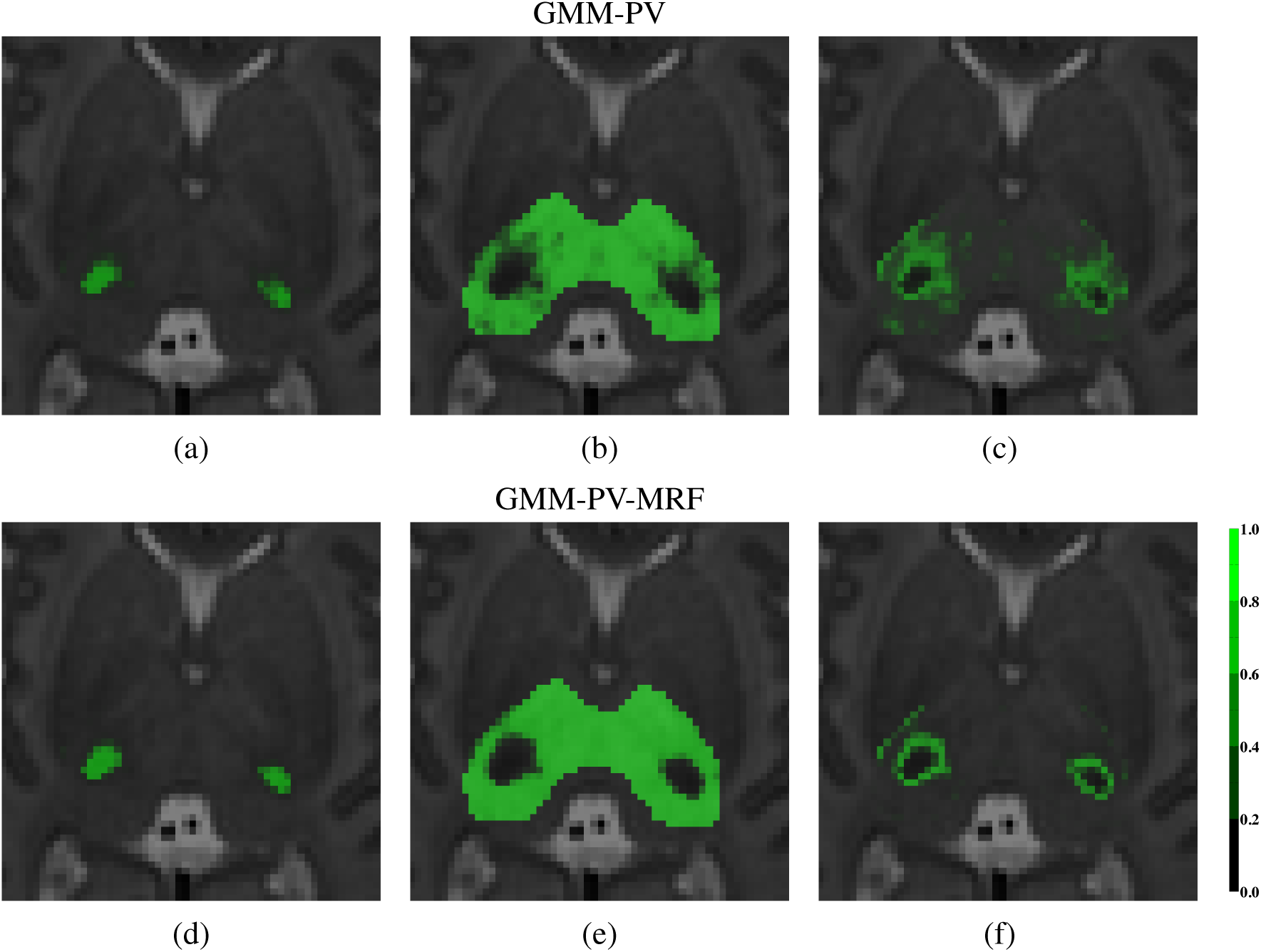
Posterior probability maps (PPMs) estimated using GMM-PV and GMM-PV-MRF (see Table 1 for method descriptions) in the thalami of a subject at 37 gestational weeks. The columns from left to right show the PPMs in the axial view for myelin-like signals (MLS), background (BKG) and partial volume voxels (PV). The PV class in GMM-PV spreads across the thalami due to the absence of spatial regularization. In contrast, GMM-PV-MRF configures spatial dependencies of all the classes via second-order MRFs and indicates the PV locations accurately at the boundary between MLS and BKG.

**Figure 9:**
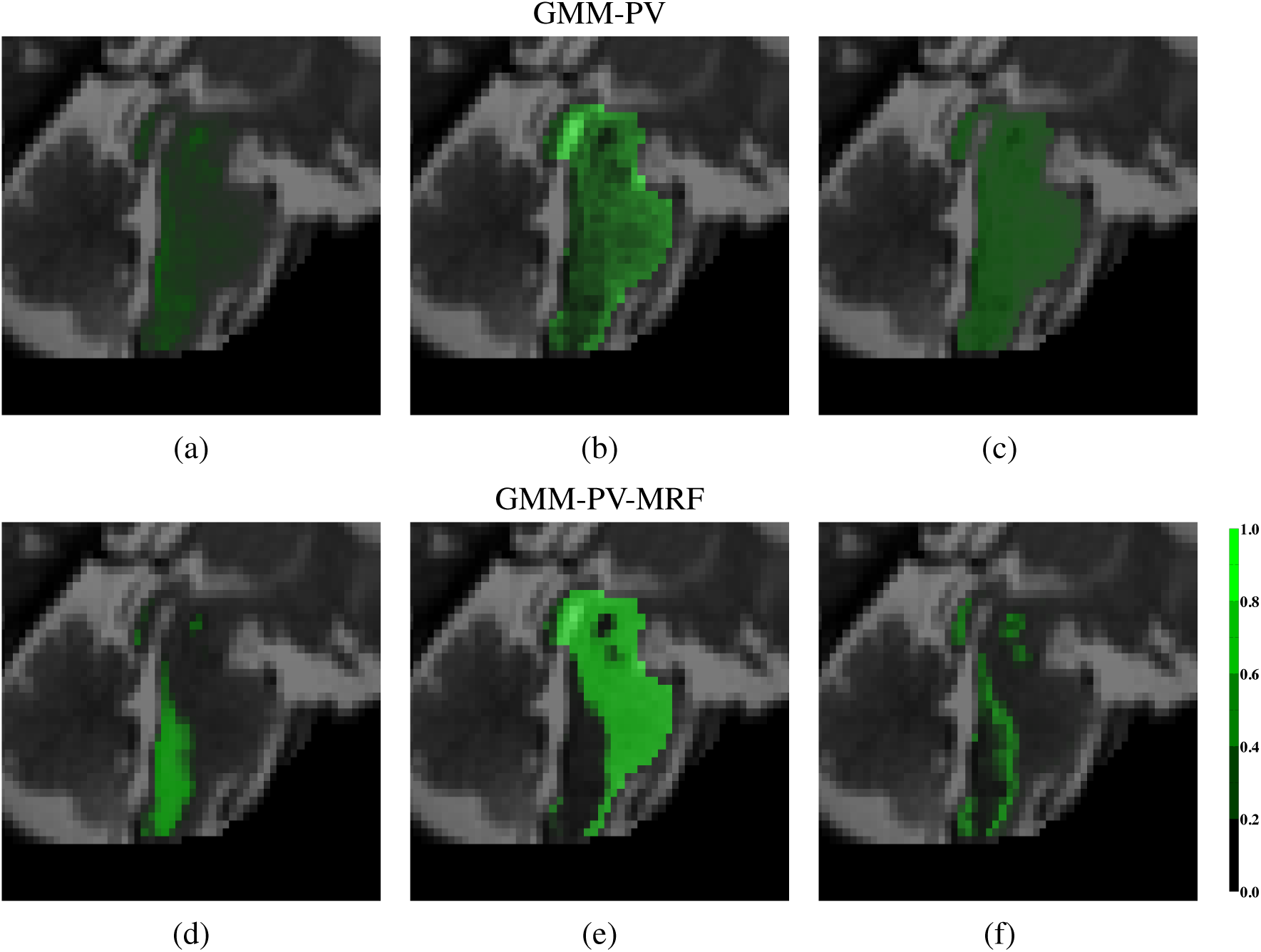
Posterior probability maps (PPMs) estimated using GMM-PV and GMM-PV-MRF (see Table 1 for method descriptions) in the brainstem of a subject at 37 gestational weeks. The columns from left to right show the PPMs in the sagittal view for myelin-like signals (MLS), background (BKG) and partial volume voxels (PV). GMM-PV-MRF performs accurately and consistently as in the thalami, whereas PV estimation using GMM-PV becomes much worse than in the thalami and shows little variation in space. This is consistent with the nearly overlaid Gaussians of the BKG and PV classes in Fig. 7(b).

### 4.3. Dependence on the initial threshold

We initialized MLS segmentations for all the methods in Table 1 using an intensity percentile threshold estimated from the volume fraction of manually annotated MLS in the ROI. However, manual reference is usually unavailable in practice, without which it is difficult to estimate the MLS content as it can vary with many factors, such as brain regions and GAs. Therefore, we continued to investigate the dependence of each method on the initial threshold.

We segmented MLS in the thalami and brainstem for the 16 test subjects by thresholding, GMM, GMM-PV and GMM-PV-MRF based on a range of initial thresholds. At each threshold, we calculated the average DC of the resultant segmentations over the subjects for each method, which was then plotted against the threshold in Figs. 10 and 11.

**Figure 10:**
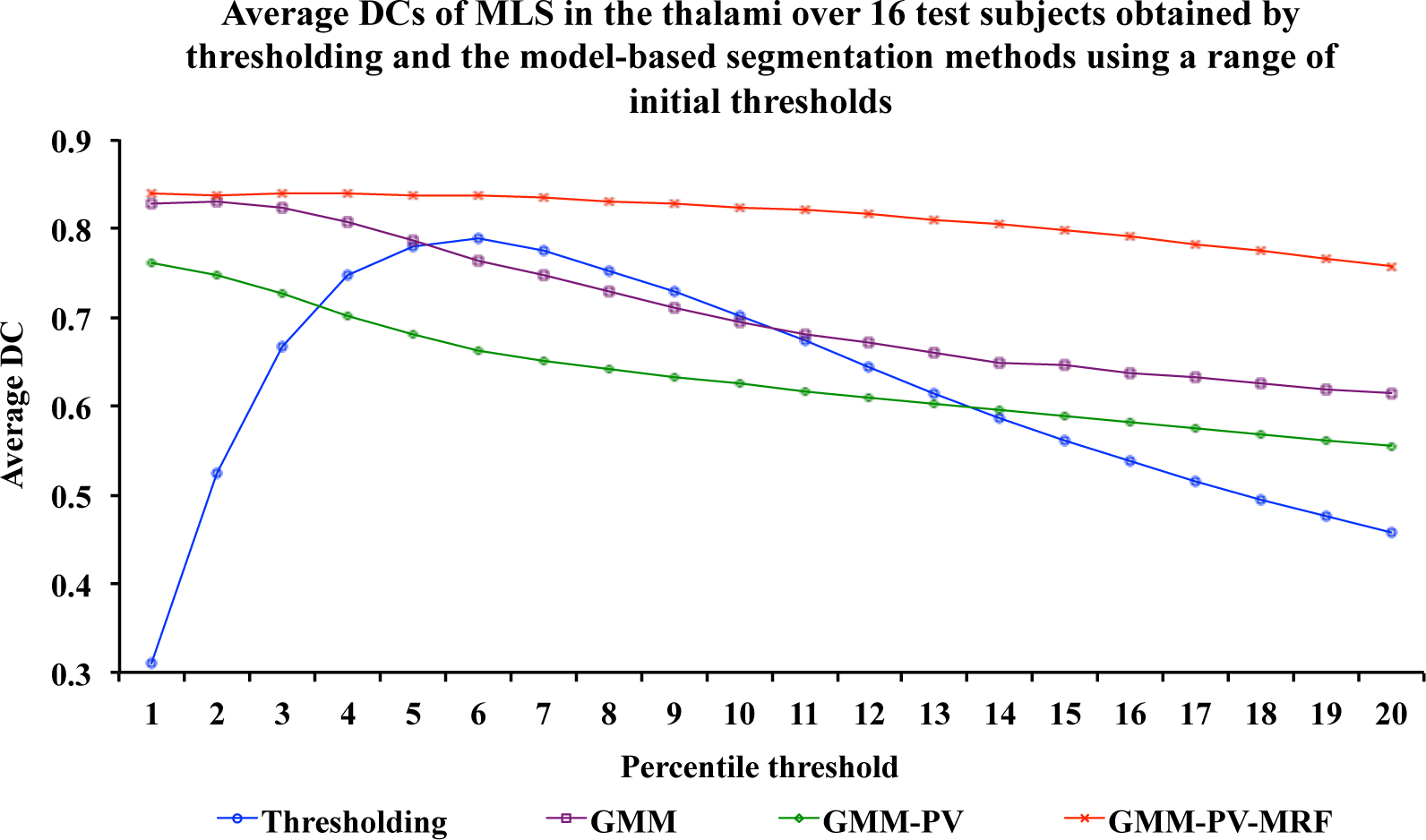
Average Dice coefficients (DCs) of myelin-like signals (MLS) in the thalami over 16 test subjects obtained by thresholding and the model-based methods (see Table 1 for method descriptions) using a range of initial thresholds. GMM-PV-MRF performs most robustly with the highest DCs as the threshold varies from the 1st to 20th percentile of the T_2_w intensities in this region. Thresholding shows considerable changes of the DC values, and the additional Gaussian in GMM-PV lowers the DCs compared to GMM due to the absence of spatial regularization.

**Figure 11:**
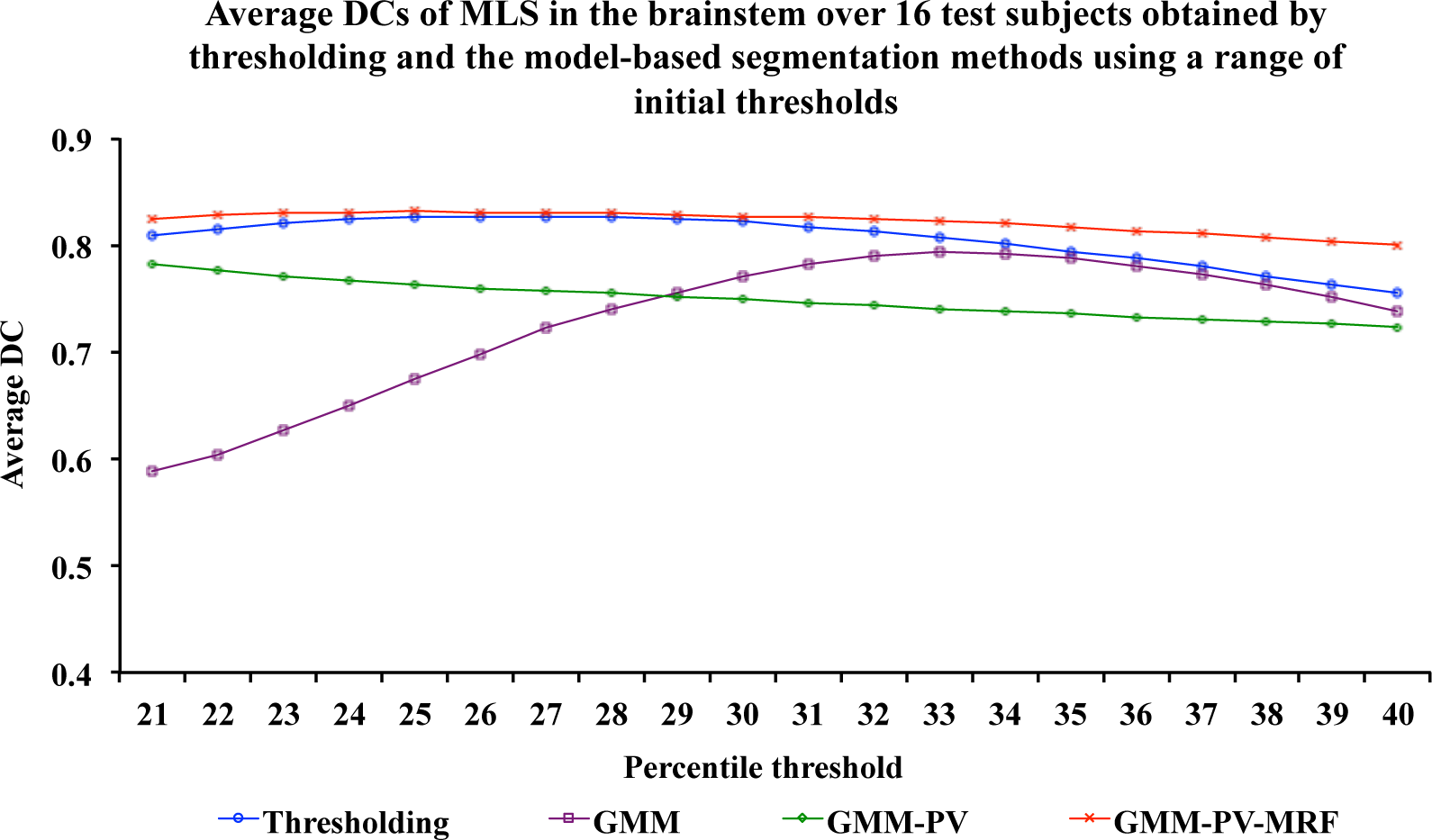
Average Dice coefficients (DCs) of myelin-like signals (MLS) in the brainstem over 16 test subjects obtained by thresholding and the model-based methods (see Table 1 for method descriptions) using a range of initial thresholds. GMM-PV-MRF performs most stably and most accurately with nearly constant DCs throughout the threshold range from the 21st to 40th percentile. The DCs of thresholding are much less affected by the varying threshold because the brainstem contains more myelinated tissue than the thalami. GMM-PV helps to stabilize the variations shown by GMM in the presence of substantial partial volume voxels, but nonetheless lowers the DCs of the baseline initial segmentations obtained from thresholding.

GMM-PV-MRF performed most stably and most accurately as the initial threshold varied from the 1st to 20th percentile in the thalami, and from the 21st to 40th percentile in the brainstem. The segmentation robustness towards the threshold is a significant advantage as the choice of the initial threshold is difficult without manual annotations, and moreover, the need for choosing the initial threshold is undesirable for practical applications. Especially when comparing the automatic MLS segmentation of an individual subject against the average growth trajectory for assessing brain maturation, it is of great importance that the volume fraction of the initial MLS estimate does not influence the final segmentation.

Thresholding, GMM and GMM-PV were particularly sensitive to the initial threshold, and performed inconsistently in different ROIs. Compared to the brainstem, the DCs obtained by thresholding underwent considerable changes in the thalami. The additional Gaussian in GMM-PV lowered the DCs of GMM in the thalami due to the absence of spatial regularization. In the brainstem with substantial PV voxels, GMM-PV helped to stabilize the variations shown by GMM, but was nonetheless unable to outperform the baseline method of thresholding.

Lastly, we verified that the 6th and 25th percentiles of the T_2_w intensities in the thalami and brainstem, respectively, optimized the DCs that could possibly be achieved using a single threshold for all the 16 subjects. Nevertheless, even the optimal threshold only performed equally well as GMM-PV-MRF in the brainstem, and could not reach the DC level of GMM-PV-MRF in the thalami.

### 4.4. Comparison to the intra-rater reliability

We assessed the intra-rater reliability as DCs between two sets of manual annotations that were available for eight of the 16 test subjects at two-week intervals between 30 and 44 weeks GA. We evaluated the automated model-based methods with respect to each manual set. The average DCs over the eight subjects are shown in Table 4 along with the *p*-values of two-tailed Student’s *t*-tests with the intra-rater DCs at the 5% significance level. It can be seen that only GMM-PVMRF performed equally well as a human rater in both ROIs, and demonstrated no significant differences with respect to the repeated manual annotations.

**Table 4:**
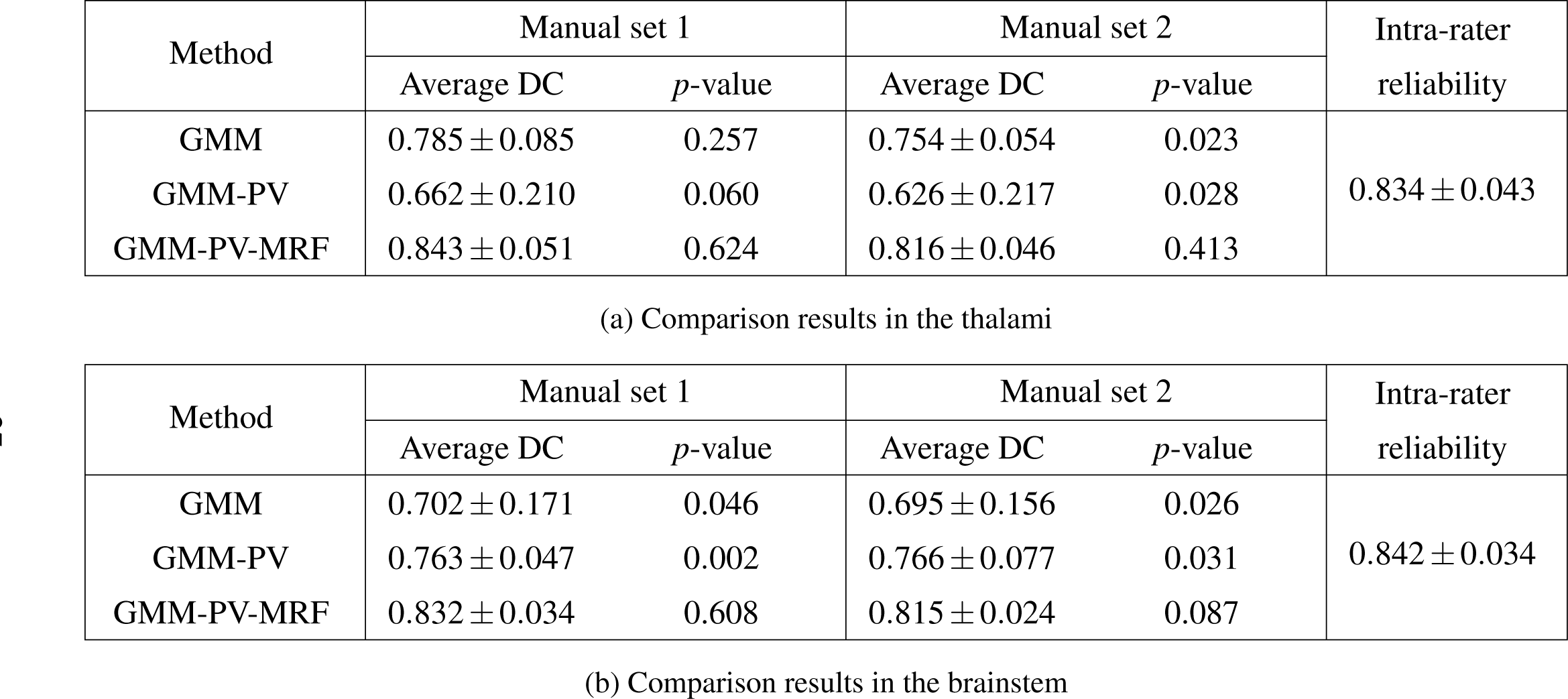
Average Dice coefficients (DCs) (*±* standard deviations) over eight test subjects obtained by thresholding and the model-based segmentation methods (see Table 1 for method descriptions) with respect to two sets of manual annotations compared to the intra-rater reliability. The *p*-values indicate the outcomes of two-tailed Student’s *t*-tests with the intra-rater DCs at the 5% significance level. The performance of GMM-PV-MRF is close to the intra-rater reliability in both regions of interest.

## 5. Applications

### 5.1. Volume growths of MLS in the ROIs

Based on the results in Section 4, we applied GMM-PV-MRF as proposed to segment MLS in the ROIs for 114 preterm infants aged between 29 and 44 weeks GA. The volumes of the resultant segmentations in the thalami and brainstem were plotted against GA in Figs. 12 and 13 respectively.

We found that the MLS volume appeared to grow exponentially with GA in the thalami, and this trend persisted in the MLS volume fraction after we corrected for the different thalamus sizes in individual subjects. In contrast, the MLS volume appeared to grow linearly with GA in the brainstem, which resulted in an approximately constant MLS volume fraction after correcting for the different brainstem sizes in individual subjects. This is because most of the brainstem appears to have completed myelination before 29 weeks GA, whereas new brain structures become myelinated in the thalami between 29 and 44 weeks GA [7, 8, 28]. Consequently, the MLS volume increased in the brainstem mainly due to the overall brain growth, and the increase in the thalami was attributed to both brain growth and appearance of new myelination. In the next section, we demonstrate the different myelination progressions in the ROIs by constructing spatio-temporal growth models for MLS.

**Figure 12:**
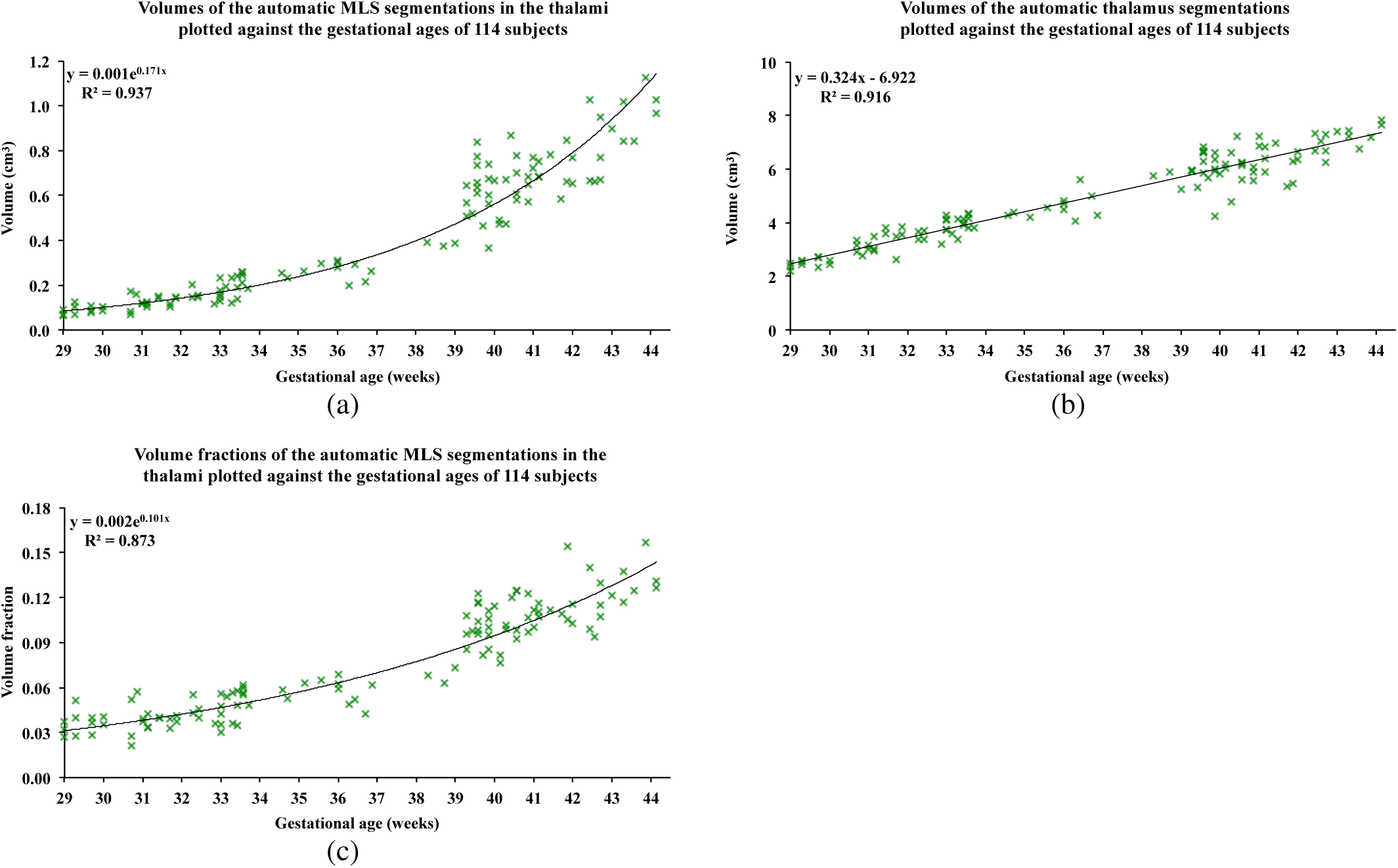
(a) Volumes of the automatic segmentations for myelin-like signals (MLS) in the thalami obtained using the proposed method GMM-PV-MRF, (b) volumes of the automatic thalamus segmentations obtained during image preprocessing, and (c) volume fractions of MLS plotted against the gestational ages (GAs) of 114 subjects between 29 and 44 weeks GA. The MLS volume appears to grow exponentially with GA, and this trend persists in the MLS volume fraction after correcting for the different sizes of the thalami in individual subjects.

**Figure 13:**
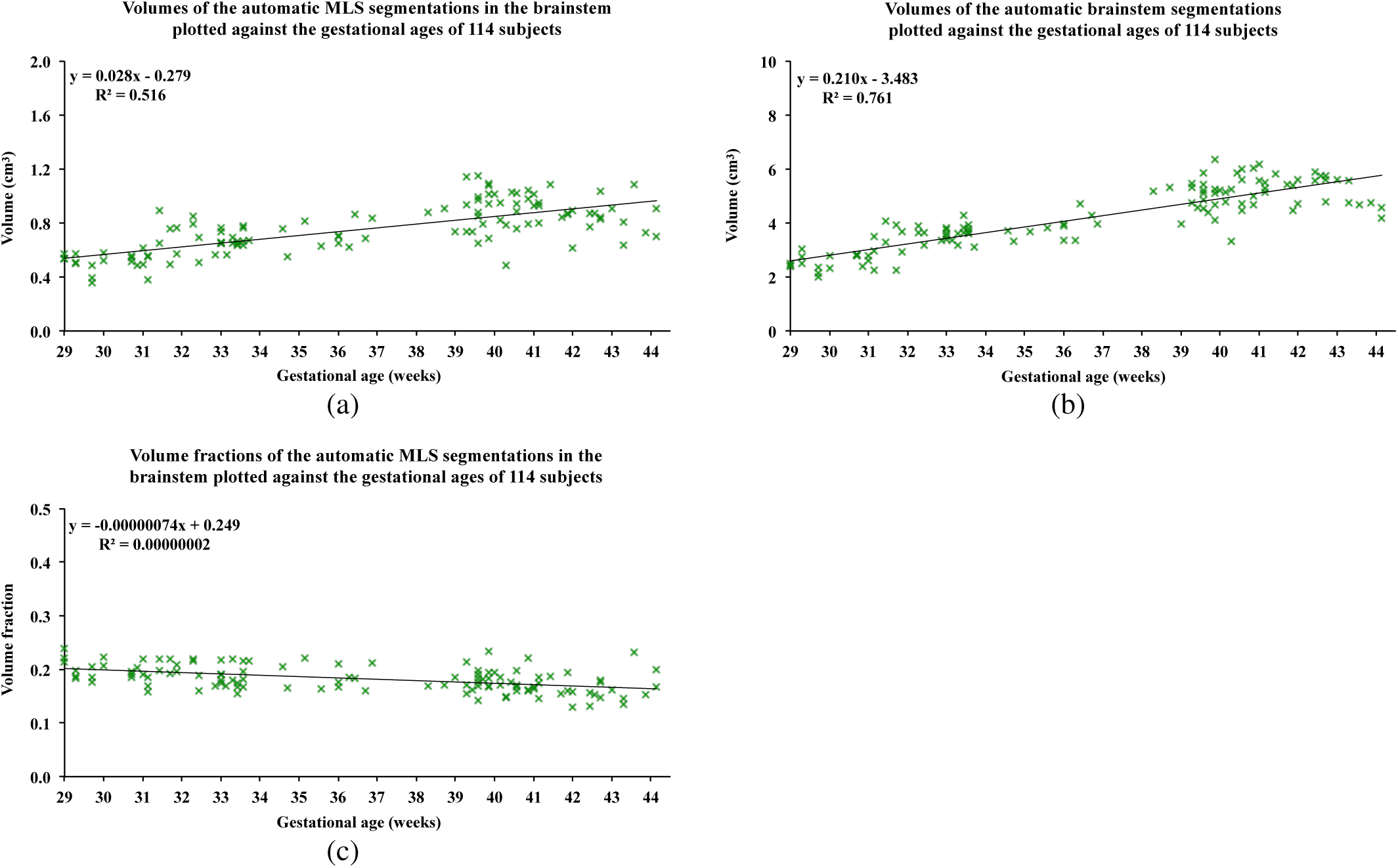
(a) Volumes of the automatic segmentations for myelin-like signals (MLS) in the brainstem obtained using the proposed method GMM-PV-MRF, (b) volumes of the automatic brainstem segmentations obtained during image preprocessing, and (c) volume fractions of MLS plotted against the gestational ages (GAs) of 114 subjects between 29 and 44 weeks GA. The MLS volume appears to grow linearly with GA, which results in an approximately constant MLS volume fraction after correcting for the different sizes of the brainstem in individual subjects.

### 5.2. Spatio-temporal growth models for MLS in the ROIs

In order to assess brain development in preterm neonates, we constructed MLS growth models in the thalami and brainstem using voxelwise logistic regression based on the automatic segmentations computed using GMM-PV-MRF for the 114 subjects. This part of our work has been published in the *Proceedings of the 13th International Symposium on Biomedical Imaging* (ISBI) [37].

First we registered the T_2_w image of each individual subject with the dilated ROI of the single reference subject at 36 weeks GA using FFD non-rigid registration [27]. This was the same reference subject used to create the ROI masks of the thalami in Section 3.1. We used normalized mutual information (NMI) as the similarity measure and 10 mm control point spacing. The automatic MLS segmentations were then transformed accordingly from each subject’s space to the common reference space. Lastly, we constructed the growth model in each ROI by fitting a voxelwise logistic function to the transformed segmentations.

We found that there was an evident difference in the onset and rate of myelination in the two ROIs. Fig. 14 showed that the VLN and STN in the thalami were likely to be myelinated before 29 weeks GA. Moreover, the model captured the arrival of MLS in both of the PLIC tracts at approximately 40 weeks GA, which confirmed the clinically observed time point [8]. Myelination of the PLIC tracts is an important landmark for evaluating neonatal brain development [5]. The brainstem, however, appeared to be well developed by the time of 29 weeks GA, and there was relatively little new myelination in the age range of our investigation. Note that because we corrected for the different sizes of each ROI in individual subjects through non-rigid registration [27], the simulated MLS progression was entirely due to the appearance of new myelinated brain structures without contribution from the overall brain growth.

**Figure 14:**
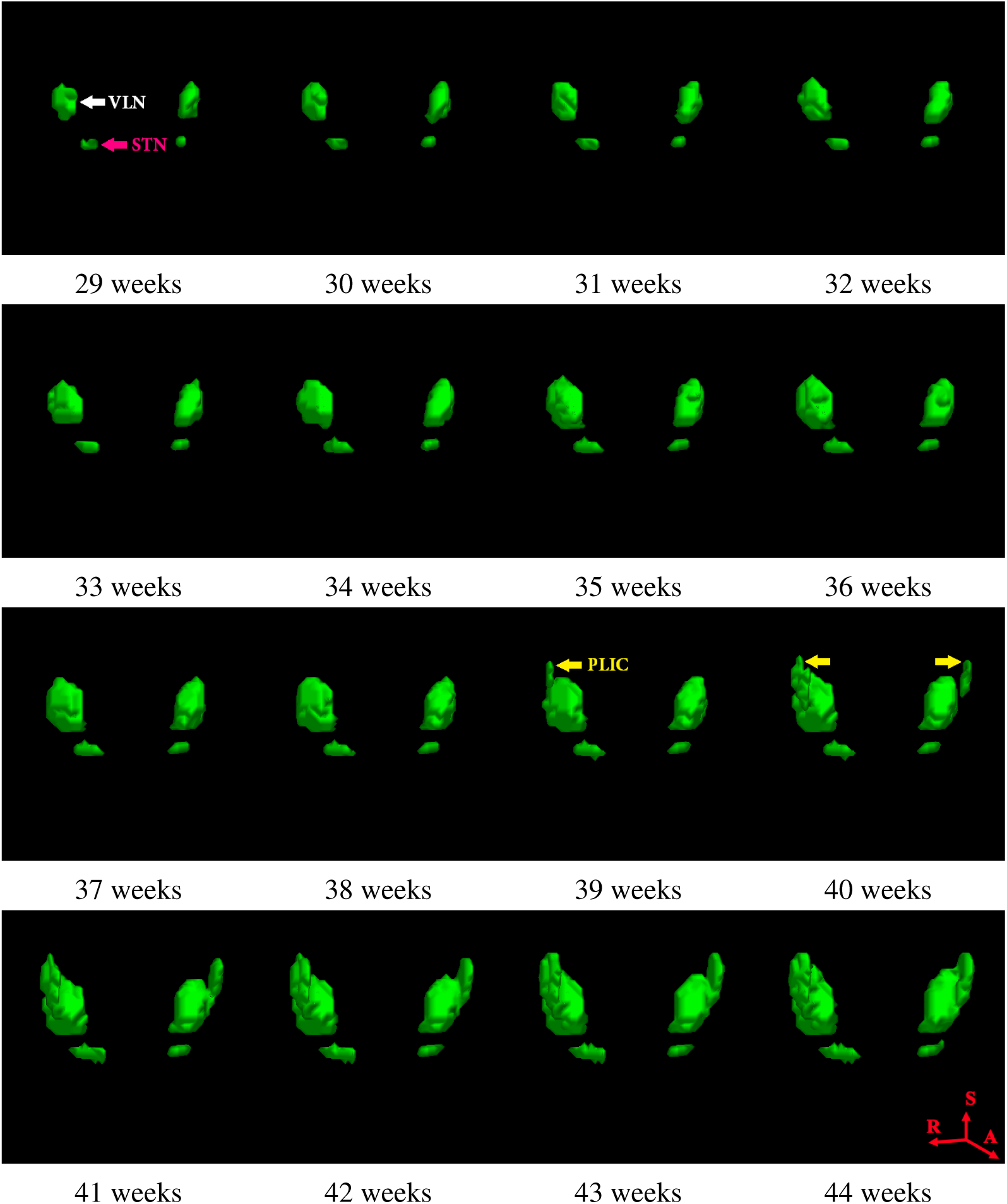
Spatio-temporal growth model for myelin-like signals (MLS) in the thalami between 29 and 44 weeks gestational age (GA). The ventrolateral nuclei (VLN) and subthalamic nuclei (STN) appear to be myelinated before 29 weeks GA. MLS becomes evident in both tracts of the posterior limbs of the internal capsule (PLIC) at approximately 40 weeks GA. This is an important landmark for evaluating neonatal brain development. As the different thalamus sizes in individual subjects have been corrected through non-rigid registration, the progression of MLS is completely due to the appearance of new myelination without contribution from the overall brain growth.

We further used the MLS growth models in the thalami and brainstem to predict GAs of the 114 preterm infants. The age estimates were determined by minimizing the sum of squared differences (SSD) between each individual transformed segmentation and the average growth model constructed in a leave-one-out procedure. The estimated GAs were plotted against the nominal values in Fig. 15. We obtained root mean squared errors (RMSEs) of 1.41 weeks and 2.56 weeks in the thalami and brainstem respectively. Therefore, each ROI has a different predictive power that best assesses a particular period of brain development. The thalami with the most prominent MLS growth between 29 and 44 weeks GA produced the most accurate age estimates for preterm infants in this age range. Details of our work in spatio-temporal modeling and age estimation can be found in [37].

**Figure 15:**
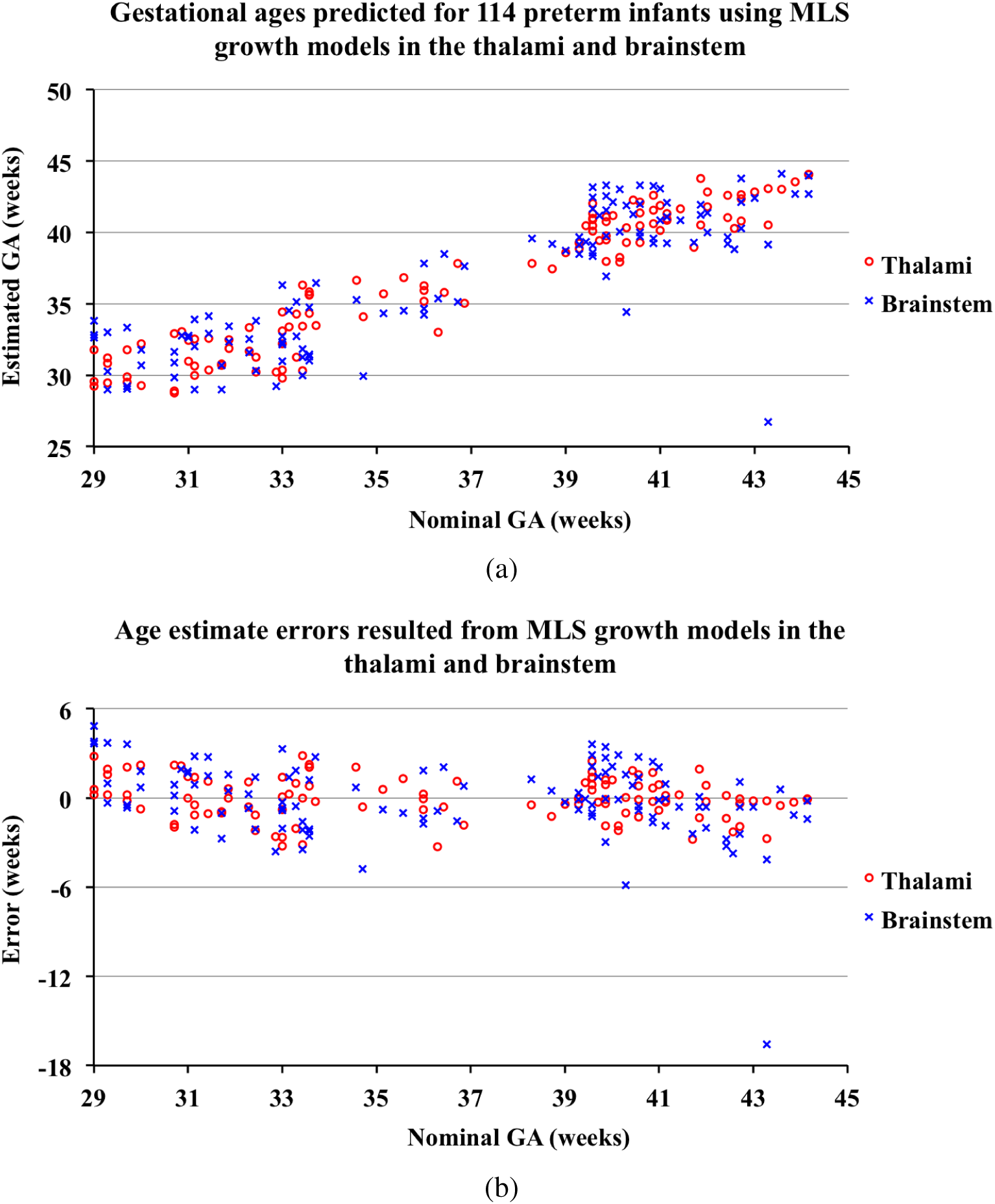
(a) Gestational ages (GAs) predicted for 114 preterm infants and (b) age estimate errors compared to the nominal GAs using the logistic growth models for myelin-like signals (MLS) in the thalami and brainstem. The predictions based on MLS in the brainstem display larger deviations from the nominal GAs than in the thalami. This is because most of the MLS spatio-temporal changes between 29 and 44 weeks GA occur in the thalami, whereas the brainstem provides much less information on progressing myelination during this particular age period.

## 6. Discussion and Conclusion

In this work, we presented an EM segmentation framework for MLS on T_2_w brain MR images of preterm neonates. The two major challenges of this task are the inadequate spatial prior information and the large number of PV voxels typical for the developing brain. The publicly available neonatal brain atlases [18, 30] do not include a probability map for myelin, and manual annotations are impractical in clinical use or for large-scale researches. The intermediate intensities of PV voxels containing both MLS and BKG make the EM optimization process unstable, and cause over-estimation of the MLS class.

To overcome these challenges, we introduced an additional Gaussian to explicitly model the PV voxels. Van Leemput *et al.* [36] observed that, without any spatial prior information, the extra Gaussian hampers segmentation robustness. The previous PV estimation methods thus used prior knowledge of the spatial locations of the composing pure tissues to guide the search for PV voxels [6, 21, 23, 31, 33, 36]. Because there are no probabilistic or manual atlases available for MLS, we identified the PV locations entirely based on tissue connectivities among the triplet of classes in the local neighborhood via second-order MRFs. This approach performed robustly and accurately with respect to manually annotated MLS in different ROIs for subjects between 29 and 44 weeks GA, regardless of the volume fraction used to obtain the initial MLS segmentations. This is crucial in practice as the choice of the initial threshold is difficult without manual annotations. The independence of the automatic MLS segmentations against the initial threshold is also important when assessing deviation from the average MLS growth model in order to estimate the brain maturation status.

In the future, we will explore multi-spectral imaging data for MLS segmentation. Myelinated GM and WM structures are shown differently using T_1_w and T_2_w sequences [4, 8]. In general, myelinated GM structures, such as the thalamic and brainstem nuclei, are more conspicuous on T_2_w images whereas myelinated WM structures, particularly the axonal tracts, are better visualized on T_1_w images [4]. Future work will include developing a multivariate EM framework that incorporates both T_1_w and T_2_w images.

In terms of the applications, we demonstrated that the proposed method for MLS segmentation provides the basis for a variety of automatic analyses that can track brain development in fragile preterm neonates using routinely acquired MR images. In particular, age estimation using the average MLS growth model can be useful in clinical practice for infants with subtle brain maturational abnormalities, who would benefit from early medical interventions soon after birth.

## Appendix

Here we derive the M-step for the proposed segmentation method (GMM-PV-MRF) where we model myelin-like signals (MLS), partial volume (PV) voxels and background (BKG) in the ROI via second-order MRFs.

We assume that the observed intensities ***y*** are generated by the hidden labels ***z*** with Gaussian probability density functions. The Gaussian parameters Φ*_y_* consist of mean *μ_k_* and standard deviation (SD) *σ_k_* of class *k* that are updated in the M-step. The hidden labels are assumed to be a realization of the random process characterized by the MRF parameters Φ*_z_*.

Denoting Φ = {Φ*_y_,* Φ*_z_*}, Van Leemput *et al.* [35] showed that the conditional expectation of the log likelihood for the complete data *{****y, z****}* can be written as:

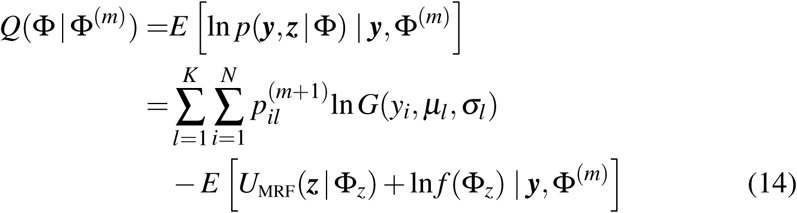

where

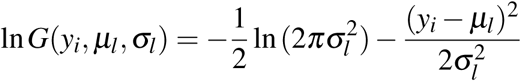

The energy function *U*_MRF_(***z*|**Φ*_z_*) is computed using Eq. 10 and *f*(Φ*_z_*) denotes a normalization constant summed over all possible configurations of the hidden labels. Explanations of the other symbols can be found in Section 3.2. Throughout the EM iterations, the MRF parameters Φ*_z_* remain constant as assigned in the connectivity tensor (Fig. 5), and hence do not contribute to the optimization of the Gaussian parameters Φ*_y_* which can be estimated in the same way as a GMM with independent voxels.

We approximate the PV class mean as the arithmetic mean of the composing tissue means (Eq. 5) and assume that the SDs of the MLS, PV and BKG classes are identical (Eq. 6). Maximization of *Q*(Φ|Φ(^*m*^)) with respect to 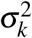 by substituting Eq. 6 into Eq. 14 yields Eq. 3 for updating the identical SD. We then differentiate *Q*(Φ|Φ(^*m*^)) with respect to each of the composing tissue means by substituting Eq. 5 into Eq. 14, and obtain the following system of linear equations:

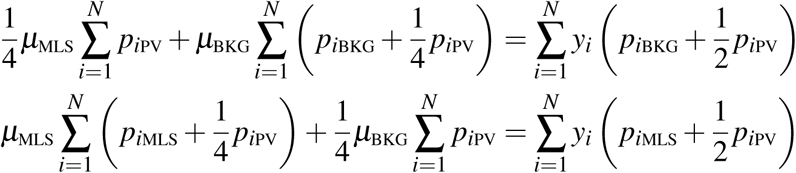

We omit the iteration number in the posterior probabilities for clarity. Lastly, we solve the linear equations via matrix inversion, as shown in Eq. 7.

## Acknowledgement

S. Wang is a recipient of the Oxford Clarendon Fund Scholarship which is provided by the Oxford University Press through the Life Sciences Interface Doctoral Training Center. The authors would like to thank the Center for the Developing Brain, King’s College London, UK, for sharing the MRI data used in this study.

## Footnotes

2 http://www.fil.ion.ucl.ac.uk/spm

3 http://brain-development.org/brain-atlases

